# Integrative structural Investigation on the architecture of human Importin4_histone H3/H4_Asf1a complex and its histone H3 tail binding

**DOI:** 10.1101/181321

**Authors:** Jungmin Yoon, Seung Joong Kim, Sojin An, Alexander Leitner, Taeyang Jung, Ruedi Aebersold, Hans Hebert, Uhn-Soo Cho, Ji-Joon Song

**Author notes:** **Abbreviations** NLS, Nuclear Localization Sequence; EM, Electromicroscopy; XL-MS, Crosslinking Massspectrometry; SAXS, Small Angle X-ray scattering; DSS, Disuccinimidyl suberate.

## Abstract

Importin4 transports histone H3/H4 in complex with Asf1a to the nucleus for chromatin assembly. Importin4 recognizes the nuclear localization sequence located at the N-terminal tail of histones. Here, we analyzed the structures and interactions of human Importin4, histones and Asf1a by cross-linking mass spectrometry, X-ray crystallography, negative-stain electron microscopy, small-angle X-ray scattering and integrative modeling. The XL-MS data showed that the C-terminal region of Importin4 interacts extensively with the histone H3 tail. We determined the crystal structure of the C-terminal region of Importin4 bound to the histone H3 peptide, thus revealing that the acidic path in Importin4 accommodates the histone H3 tail and that histone H3 Lys14 is the primary residue interacting with Importin4. Furthermore, the molecular architecture of the Importin4_histone H3/H4_Asf1a complex was produced through an integrative modeling approach. Overall, this work provides structural insights into how Importin4 recognizes histones and their chaperone complex.

## Introduction

The nucleosome is the basic unit of chromatin and consists of approximately 146 bp of DNA wrapped around a histone octamer [1]. The dynamic structure of chromatin and its regulations play critical roles in various cellular processes such as DNA replication, transcription and repairs [2]. During DNA replication, nucleosomes are actively assembled into duplicated DNA to maintain the integrity of the chromatin structure. To support this process, a large number of histones need to be synthesized and transported into the nucleus. Previous studies have described the processes by which the newly synthesized histones H3 and H4 are deposited onto DNA [3–5]. Newly synthesized histone H3/H4 dimers are escorted by several histone chaperones including Nuclear Autoantigenic Sperm Protein (NASP), Histone Acetyltransferase 1 (HAT1), RbAp48 and Anti-silencing factor 1a (Asf1a), Chromatin Assembly Factor-1 (CAF-1) and Regulator of Ty1 Transposition (Rtt106). Asf1a, the major histone H3/H4 chaperone, binds to the tetramer interface between histone H3/H4 dimers, thereby preventing these dimers from binding to other proteins inappropriately and blocking tetramer formation before nucleosome assembly. The histone H3/H4_Asf1a complex is then transported by Importin4 to other histone chaperones, such as the CAF-1 complex, in the nucleus [3, 4, 6–8]. H2A and H2B have their own histone chaperones such as Nucleosome assembly protein 1 (Nap1) [9]. In addition, Kap114p in yeast is responsible for delivering the histones H2A/H2B into the nucleus [10].

Importin4 is a major histone H3/H4 nuclear import protein in humans and is a member of the Karyopherin-β superfamily. The proteins in this family have low sequence homology to one another (below 20%) and consist of tandemly arrayed HEAT repeats which promote flexibility and elongated conformations of the structures [11–13]. These features allow one protein to accommodate several different cargoes in addition to directly recognizing the nuclear localization sequences (NLSs) of the cargoes. Importin4 is also responsible for transporting other cargoes in addition to histone H3/H4, including the ribosomal protein S3a, the vitamin D receptor, hypoxia inducible factor-α and human epididymis protein 4 [14–16]. Several previous studies showed that other nuclear importins are also able to transport histone H3 [4, 17–22], and recent biochemical and structural studies have shown that other nuclear importins are also able to transport histone H3 [23, 24]. However, Importin4 is the *bona fide* nuclear importin for histone H3 and H4 [3, 4]. In yeast, Kap123 is a homolog of human Importin4 and recognizes the NLSs residing in the N-terminal tails of both histone H3 and H4. Previous studies have defined the minimal NLSs of histones H3 and H4 as amino acid (a.a.) residue 1-28 of histone H3 and 1-21 a.a. of histone H4 for yeast Kap123, and have shown that these NLSs are also critical for the interaction with human Importin4 [10, 19]. In addition, recent biochemical study mapped the regions in histone H3 and H4 tails important for Importin4 binding [23]. Although Importin4 plays a major role in delivering new histone H3/H4_Asf1a complexes to the nucleus in humans, little is known about the molecular architecture of the Importin4_histone H3/H4_Asf1a complex and the mechanism by which Importin4 interacts with the histone NLS.

In this study, we investigated the structure and molecular interactions of the Importin4_histone H3/H4_Asf1a complex through an integrative modeling approach relying on multiple structural and proteomic data sources, including cross-linking mass-spectrometry (XL-MS), X-ray crystallography, small angle X-ray scattering (SAXS), and negative-stain single-particle electron microscopy (EM). The resulting integrative model of the complex together with the experimental data enables us to understand the mechanism by which Importin4 recognizes the histone H3/H4_Asf1a complex.

## Result

### Cross-linking Mass Spectrometry Analysis of the Importin4_histone H3/H4-Asf1 Complex

Importin4 is the major nuclear importin for newly synthesized histone H3/H4 complexed with the histone chaperone Asf1a. To investigate the interactions among these proteins, we performed XL-MS on the Importin4_histone H3/H4_Asf1a complex. The highly purified complex was cross-linked with disuccinimidyl suberate [25] and the homogeneity of the cross-linked sample was analyzed by native PAGE (Fig. 1a). The cross-linked sample was then subjected to XL-MS analysis (Supplementary Table). To investigate the interactions among Importin4, histone H3/H4 and Asf1a, we mapped the cross-linked residues on the crystal structure of the histone H3/H4_Asf1a complex (PDB ID: 2IO5). Most of the cross-linked residues in the histones are located on the DNA binding surface, thus revealing that Importin4 probably recognizes the DNA binding surface of histone H3/H4 (Fig. 1c and d).

**Fig. 1.**
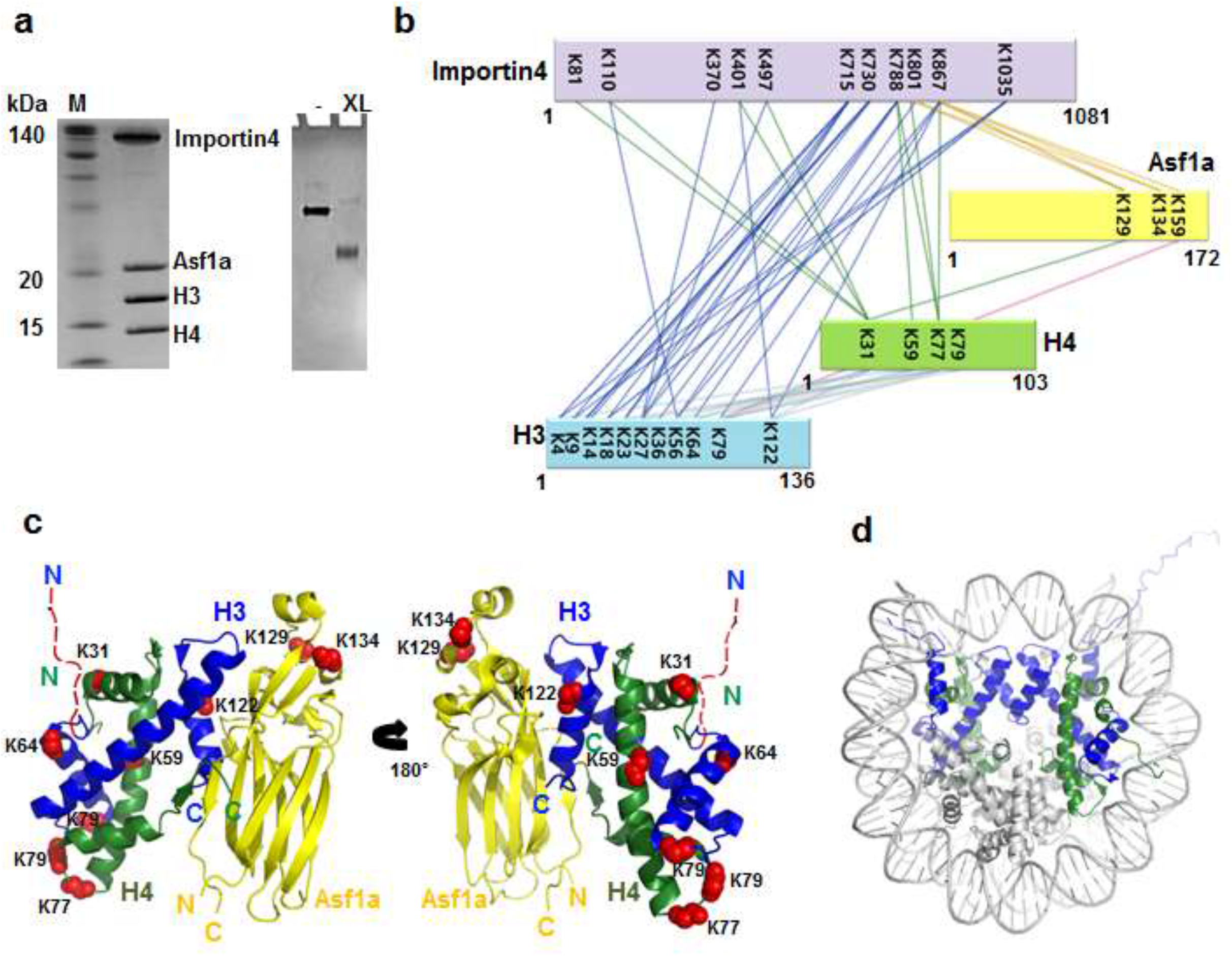
Cross-linking mass spectrometry analysis of the Importin4_histoneH3/H4_Asf1a complex. **(a)** Purified Importin4_histone H3/H4_Asf1a complex analyzed by SDS-PAGE (left panel). The DSS cross-linked complex analyzed by native PAGE (right panel) shows the homogeneity of the samples. The cross-linked protein migrated faster than the uncross-linked protein due to its compactness on a native PAGE. **(b)** The cross-linked residues are linked with dotted lines on the primary structures of Importin4 (pale purple), histone H3 (cyan), histone H4 (green) and Asf1a (yellow). The list of the cross-linked residues is shown in Supplemental Table 1. **(c)** The residues of histone H3/H4_Asf1a (PDB ID: 2IO5) that were cross-linked with Importin4, are shown in space filling model in red in two different orientations, thus showing that Importin4 might bind the DNA binding surface of the histone H3/H4 dimer. The red dashed line indicates the disordered N-terminal tail of histone H3. **(d)** The nucleosome structure (PDB ID: 1AOI) with histone H3 in blue and histone H4 in green shows the DNA binding surface of histone H3/H4.

It is notable that we also observed cross-links between Asf1a and Importin4, thus suggesting that Asf1a might contribute to the Importin4 binding.

Furthermore, our XL-MS data showed that the N-terminal tail of histone H3 interacts extensively with the C-terminal region of Importin4, whereas there is little cross-linking between Importin4 and the N-terminal tail of histone H4, despite of the presence of several lysine residues in the histone H4 tail (Fig. 1b). Almost all of the lysines (K4, K9, K14, K18, K23, K27) in the histone H3 tail are extensively crosslinked with the region from Lys715 to Lys1035 of Importin4, thus indicating that this region accommodates the N-terminal tail of histone H3. In the case of histone H4, the lysines K31, K59 and K77 located at the DNA binding surface in the nucleosome structure were cross-linked with Importin4. We also observed that Importin4 is crosslinked with the histone fold of the H3/H4 dimer and Asf1a, suggesting that not only the tail, but also the histone fold as well as Asf1a might contribute to the Importin4 binding. Consistent with this, the previous work by Soniat et al also reached similar conclusion by biochemical analysis [23]. These data delineate the interactions among Importin4, histone H3/H4 and Asf1a, showing that the N-terminal tail of histone H3 predominantly interacts with the C-terminal region of Importin4 and that Importin4 masks the DNA binding surface of histone H3/H4.

### Importin4 interacts with the N-terminal tail of histone H3

It was shown that the NLSs located at the N-terminal tails of histone H3/H4 play critical roles in translocation of the histone to the nucleus [10, 19]. XL-MS data indicates that the N-terminal tail of histone H3 was extensively crosslinked with Importin4. Therefore, we hypothesized that Importin4 directly interact with the N-terminal tail of histone H3 in the Importin4_histone H3/H4_Asf1 complex. We first performed a limited-proteolysis using trypsin on Importin4_histone H3/H4_Asf1 complex and histone H3/H4_Asf1 complex to examine whether the N-terminal tail of histone H3 is protected upon Importin binding to H3/H4_Asf1. In agreement with XL-MS data, our limited proteolysis experiments showed that the N-terminal tail of histone H3 but not histone H4 was protected from tryptic digestion in the presence of Importin4, thus further confirming that Importin4 primarily interacts with the N-terminal tail of histone H3 (Fig. 2a). We then examined the direct binding with the N-terminal tail of histone H3 and the C-terminal region of Importin 4 (C-Importin4, 668 to 1081 a.a.) having the extensive crosslinking between histone H3 tail by Surface Plasmon Resonance (SPR). We immobilized C-Importin4 on the CM5 chip and measured the bindings between C-Imporin4 and several N-terminal regions of histone peptides (Fig. 2b, 2c and 2d). The SPR data show that the histone peptide (1-18 a.a.) binds to C-Importin4 whereas other peptides (1-11 a.a. and 1-6 a.a.) did not bind, suggesting that N-terminal histone H3 which contained _12_GGKAPRK_18_ residues interacts with C-Importin4.

**Fig. 2.**
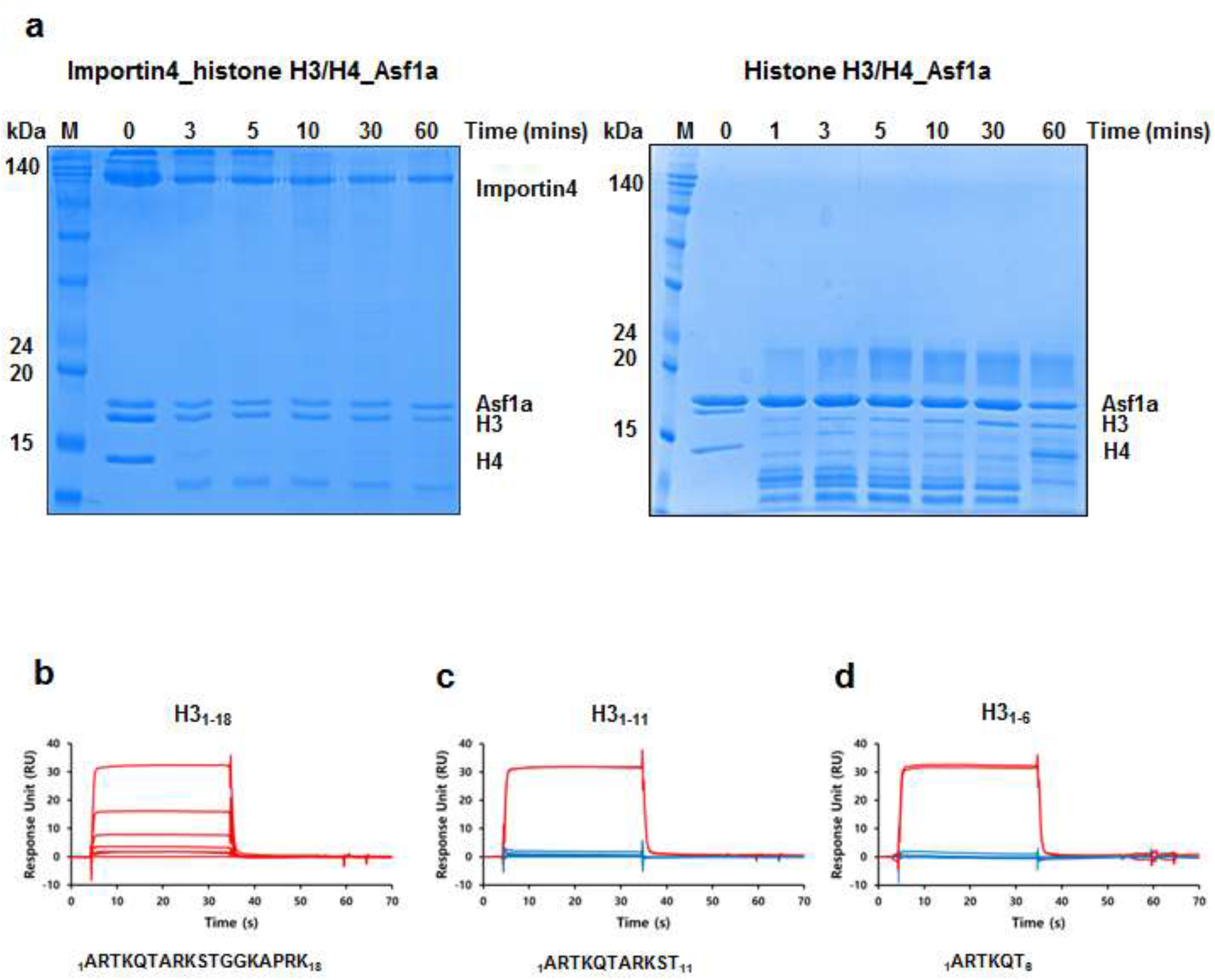
Limited proteolysis Analysis of the Importin4_histone H3/H4_Asf1 Complex. **(a)** Limited proteolysis of Importin4_histone H3/H4_Asf1a and histone_H3/H4_Asf1a. Importin4_histone H3/H4_Asf1a (left panel) or histone H3/H4_Asf1a (right panel) were incubated with trypsin for 60 min at 37 °C. The reaction products were analyzed on a 15% SDS-PAGE and stained with Coomassie brilliant blue staining. **(b-d)** Interactions between C-Importin4 and deletion histone H3 peptides (H3_1-18_: 1 to 18 a.a, H3_1-11_: 1 to 11 a.a. and H3_1-6:_ 1 to 6 a.a.). The C-Importin4 was immobilized with 6,000 response units (RU) on a CM5 chip. The peptides were injected as the analytes at different concentrations (0, 0.5, 1, 2, 4, 6 and 8 μM).

### The Crystal Structure of the C-terminal Region of Importin4 and Its H3 Peptide Complex

To understand in more detail how Importin4 recognizes the histone H3 tail, we determined the crystal structure of the C-Importin4, in which the histone tails are extensively cross-linked, by using a Se-Met single anomalous diffraction (SAD) method (Table 1). The structure shows that the C-Importin4 consists of two layers of outer and inner antiparallel α-helices, similarly to other known HEAT repeat protein structures (Fig. 3a and 3b) [24, 26–28]. Helix number was given based on helices of the model of full-length Importin4 (Fig. 5a). The electrostatic surface representation of C-Importin4 shows a highly acidic patch between the H18B and H22B helices (Fig. 3c). This acidic patch might serve as a pocket for the highly positively charged NLS of histone H3. To further investigate the interaction between histone H3 and Importin4, we determined the crystal structure of C-Importin4 bound to a histone H3 peptide (1-18 a.a., _1_ARTKQTARKSTGGKAPRK_18_) through a molecular replacement method using the C-Importin4 structure as a search model (Table 1 and Fig. 2d). The structure of C-Importin4_histone H3 peptide is almost identical to that of C-Importin4 with 0.49Å RMSD indicating the histone peptide binding did not cause conformational change (Supplementary Fig. 1). Although clear extra density near the acidic patch was observed, we were able to place only four amino acids (_12_GGKA_15_) out of 18 in the electron density map (Supplementary Fig. 2). The complex structure shows that the histone H3 peptide is bound to the acidic patch located between the H18B and H22B helices of Importin4 (Fig. 3e and 3f). Specifically, the positively charged ε-amine in Lys14 of histone H3 interacts with the carboxyl groups of Importin4 Asp994 and Glu997 side chains located at the H22B helix, and Importin4 Phe921 supports the hydrocarbon of H3 Lys14 side chain, in a manner reminiscent of a histone methyltransferase binding the Lys residues of substrates. The carboxyl group of Importin4 Glu884 interacts with the carbonyl oxygen of H3 Gly13, and Importin4 Glu837 interacts with the carbonyl group of H3 Gly12. In addition, the side chain of Importin4 Asn918 supports the carbonyl oxygen of H3 Ala15 and the amine nitrogen of H3 Ala15. Furthermore, the Glu914 side chains interacts with the carbonyl oxygen of H3 Ala15 (Fig. 3f). Our structure reveals the molecular details on how Importin4 recognizes the NLS of histone H3.

**Fig. 3.**
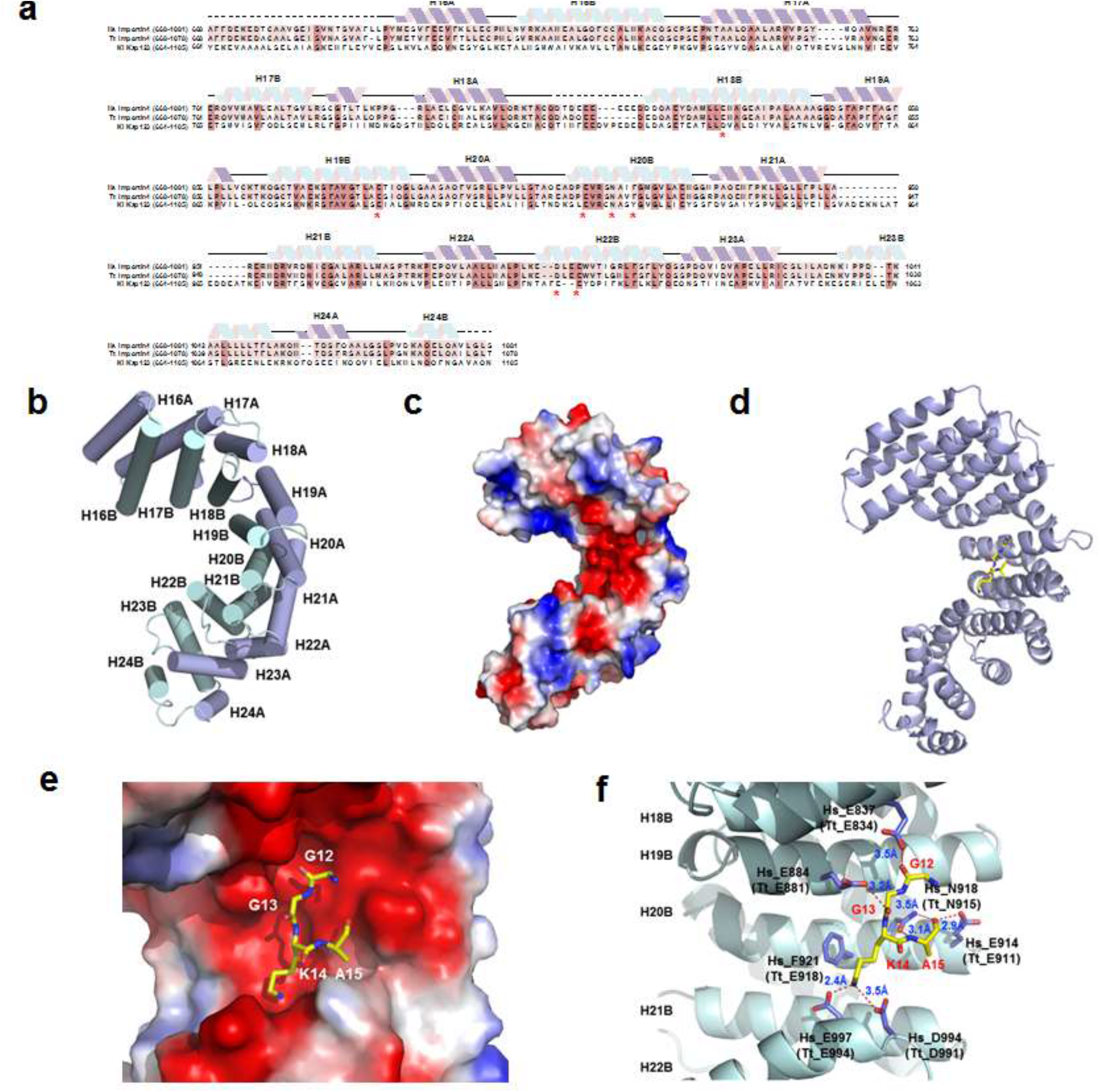
Crystal structures of the C-terminal region of Importin4 and its histone peptide complex. **(a)** Sequence alignment of the C-terminal regions of human Importin4 (*Homo sapiens*, Hs_Importin4), dolphin Importin (*Tursiops truncates*, Tt_Importin4) and yeast Kap123 (*Kluyveromyces lactis*, Kl_Kap123). The secondary structures are shown above the sequences based on the crystal structure of C-Importin4. Conserved amino acids among three Importins are shown in pink and conserved residues between human and dolphin Importin4 are shown in pale pink. Amino acids involved in binding of the histone H4 tail are indicated by asterisks. **(b)** The crystal structure of the C-terminal region (668 to1081 a.a.) of Importin4 (C-Importin4). The outer helices are labelled H16A to H24A (pale purple) and the inner helices are labelled H16B to H24B (pale cyan). **(c)** The electrostatic surface representation of C-Importin4 showing an acidic patch between the H18B and H22B helices. **(d)** The crystal structure of C-Importin4 bound to the histone H3 peptide. The histone H3 peptide (GGKA, 12-15 a.a.) is shown as a stick model in yellow. **(e)** The histone peptide (GGKA, 12-15 a.a.) is shown on the acidic patch shown in an electrostatic surface representation. **(f)** Detailed view of the histone peptide at the histone binding pocket. Lys 14 of histone H3 interacts with the carboxyl groups from Hs_E997 (Tt_E994) and Hs_D994 (Tt_D991). Importin4 Hs_E884 (Tt_E881) interacts with the the carbonyl oxygen of H3 Gly13. The carboxyl side chain of Importin4 Hs_E837 (Tt_E834) interacts with the carbonyl oxygen of H3 Gly12. The side chain of Importin4 Hs_N918 (Tt_N915) interacts with the imine nitrongen and carbony oxygen of H3 A15, which is also coordinated by Importin4 Glu914. The interactions are shown as red dotted lines with distances in Å. The residues from human Importin4 are shown as Hs and dolphin Importin4 is shown as Tt.

**Table 1.** Data collection and refinement statistics.

|  | C-Importin4<br>(SAD) | C-Importin4-H3 peptide<br>(MR) |
| --- | --- | --- |
| **Data collection** |  |  |
| Wavelength (Å) | 0.98002 | 0.97950 |
| Space group | P2<sub>1</sub> | P2<sub>1</sub> |
| a, b, c (Å) | 64.92, 110.77, 126.82 | 65.02, 108.54 124.86 |
| α = β = γ (°) | 90.00, 106.50, 90.00 | 90.00, 108.31, 90.00 |
| Resolution (Å) | 50-3.0 (3.05-3.00) * | 50-3.20 (2.36-3.20) |
| R<sub>sym</sub> | 0.10 (0.60) * | 0.12 (0.85) * |
| I / σ(I) | 34.82 (3.52) * | 24.35 (2.34) * |
| Completeness (%) | 99.2 (97.5) * | 99.4 (100.0) * |
| Redundancy | 7.3 | 4.5 |
| **Refinement** |  |  |
| Resolution (Å) | 50-3.0 | 50-3.22 |
| No. reflections | 33,882 | 26,872 |
| R<sub>work</sub> / R<sub>free</sub> | 0.226/0.280 | 0.218/0.281 |
| No. atoms |  |  |
| Protein | 11,283 | 11,165 |
| R.m.s.d. deviations |  |  |
| Bond lengths (Å) | 0.003 | 0.004 |
| Bond angles (°) | 0.739 | 0.672 |
| Ramachandran plot (%) |  |  |
| Allowed/Generously allowed | 92.9 / 7.1 | 92.1 / 7.9 |

R<sub>work</sub> : $\sum ||F_{obs}| - |F_{calc}|| / \sum |F_{obs}|$
R<sub>free</sub>: $\sum ||F_{obs}| - |F_{calc}|| / \sum |F_{obs}|$ where 10% of randomly selected data were used.
R<sub>sym</sub>: $\sum |I_{hkl} - \langle I_{hkl} \rangle| / \sum I_{hkl}$ , where $\langle I_{hkl} \rangle$ is the mean intensity of all reflections equivalent to reflection hkl.
\* The values in parentheses are for highest-resolution shell.

### Binding Analysis of Importin4 and Histone H3

To more specifically define the interactions between the histone H3 tail and Importin4, we mutagenized the residues in Importin4 that interact with the histone H3 peptide, and measured their interaction with histone H3 by Surface Plasmon Resonance (SPR). First, to examine the binding between histone H3 and C-Importin4, we immobilized a C-terminal biotinylated histone H3 peptide (1-35 a.a.) on a CM5 chip coated with streptavidin and measured the binding of C-Importin4 to the histone H3 peptide by injecting C-Importin4 at six different concentrations (0, 0.5, 1, 2, 4, 6, and 8 μM) (Fig. 4a). Fig. 3a shows that C-Importin4 bound to histone H3 peptide with micromolar KD range (Supplementary Fig. 3). These data are consistent with the XL-MS data and the crystal structure in that the histone H3 tail interacts with the C-terminal region of Importin4. The crystal structure we determined was confined to the C-terminal region of Importin4 and it is possible that other regions of Importin4 besides the C-terminal region might interact with the histone H3 tail. To examine this possibility, we used full-length Importin4 for the following experiments. However, because full-length human Importin4 alone without Histone H3/H4_Asf1a was not stable, we used human Importin4 orthologues from *Tursiops truncatus* (Dolphine, Tt_Importin4) for the SPR measurements. These proteins share more than 89% sequence identity with the human sequence (Fig. 3a and Supplementary Fig. 4). We analyzed the binding of full-length Tt_Importin4 (Tt_WT) to the immobilized histone H3 peptide by injecting full-length Tt_Importin4 at seven different concentrations (31, 62, 125, 250, 500, 1000, and 2000 nM), showing that Tt_Importin4 binds to the histone H3 peptide with slower on and off rates compared with those of C-Importin4. These data indicate that the N-terminal region beyond the C-terminal region of Importin4 could also contribute to the binding to histone H3 although it might not directly bind (Fig. 4b). We then mutagenized the residues involved in histone binding in Tt_Importin4 based on the C-Importin4 structure (Supplementary Fig. 5). Owing to the instability of the immobilized surface and the proteins during the extended time period of SPR measurements, we measured the relative binding of the mutants compared with those of the wild-type Tt_Importin4. We injected the mutants at six different concentrations (31, 62, 125, 250, 500, and 1000 nM) on the chip immobilized with histone H3 peptide (1-35 a.a.). To validate the surface stability, we injected the wild-type protein before and after each mutant measurement and used the binding of the wild-type to monitor the relative binding of the mutants to the wild-type. The crystal structure of the C-Importin4 bound histone H3 tail shows that Histone H3 Lys14 is coordinated by two carboxylate groups, Hs_D994 and Hs_D997 in human Importin4, and Tt_D991 and Tt_E994 in Tt_Importin4. To examine the contributions of these interactions, we mutagenized Tt_E994 (Hs_E997) to alanine in Tt_Importin4 and measured its binding to the histone H3 tail (Fig. 4c and 4f).

**Fig. 4.**
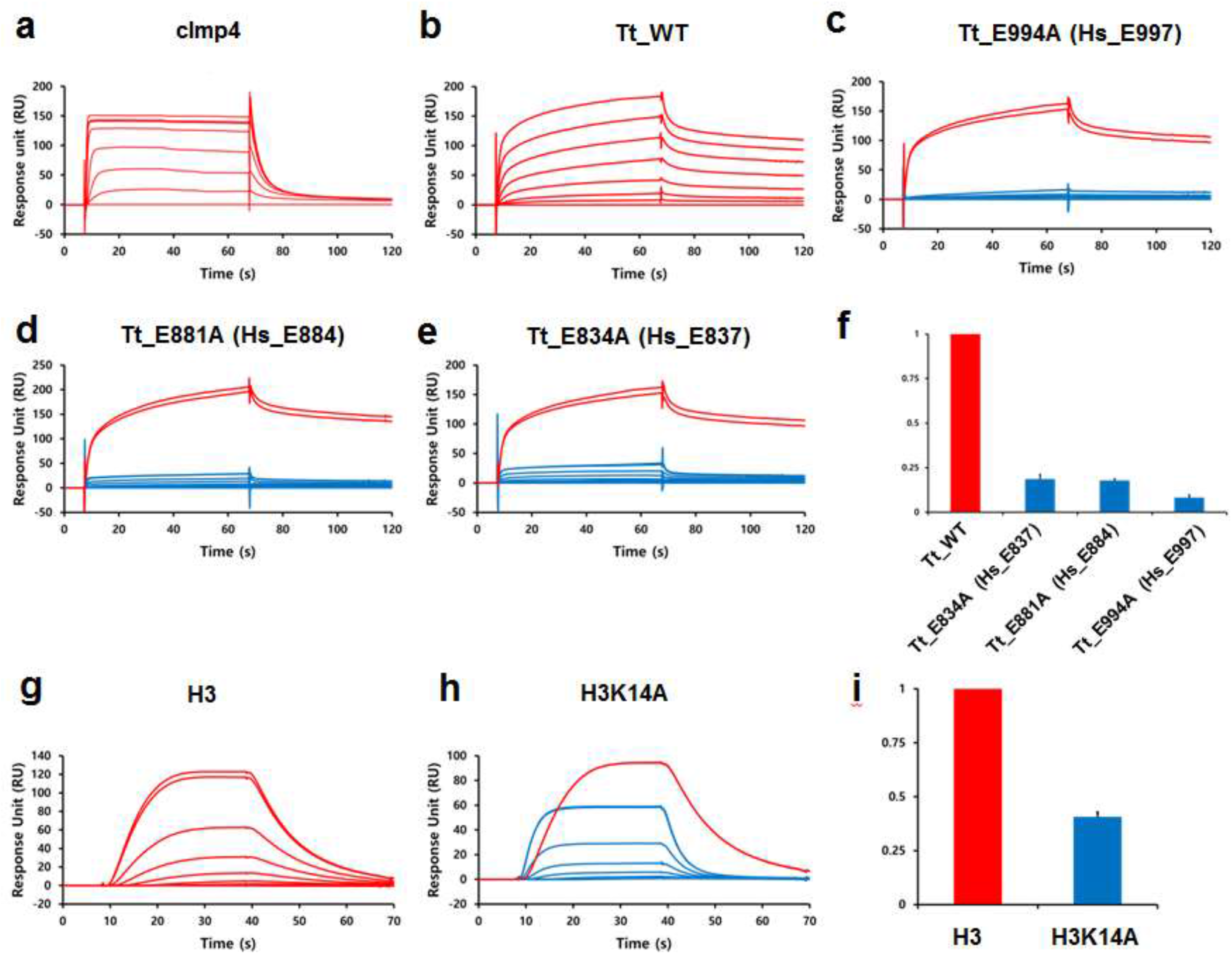
Analyzing interactions between Importin4 and NLS of histone H3 peptide by Surface Plasmon Resonance. **(a)** Interactions between C-Importin4 and a histone H3 peptide (1 to 35 a.a). The C-terminal biotinylated histone H3 peptide (1 to 35 a.a.) was immobilized with 120 response units (RU) on a CM5 chip coated with streptavidin. C-Importin4 was injected as the analyte at different concentrations (0, 0.5, 1, 2, 4, 6 and 8 μM) of C-Importin4. **(b)** Interactions between full-length Tt_Importin4 (Tt_WT) with a histone H3 peptide (1 to 35 a.a). The C-terminal biotinylated histone H3 peptide (1 to 35 a.a.) was immobilized with 120 RU on a CM5 chip coated with streptavidin. Wild-type Tt_Importin4 (Tt_WT) was injected as the analyte at different concentrations (0, 31, 62, 125, 250, 500, 1000 and 2000 nM). **(c)-(e)** The binding to histone H3 (1 to 35 a.a.) was measured for each mutant. The C-terminal biotinylated histone H3 peptide (1 to 35 a.a.) was immobilized with 120 RU on a CM5 chip coated with streptavidin. Mutants of Tt_Importin4 (C: Tt_E994A, D: Tt_E881A, and E:Tt_E834A) at different concentrations (0, 31, 62, 125, 250, 500 and 1000 nM) were injected, and the sensorgrams are shown in blue color. Tt_Importin4 (Tt_WT) at a concentration of 1,000 nM concentration was injected at before and after each injection of mutant and the sensorgrams are shown in red as controls. **(f)** The relative binding of mutants shown in (C)-(E) compared with the binding of Tt_WT is shown in a bar graph. The average of the ratios between the binding of the mutant to that of Tt_WT at four different concentrations are shown on the Y axis. The errors among the ratios were combined to generate the error bars. **(g)**-(h) The contribution of histone H3 K14 to the interaction with Importin4. Tt_Importin4 was immobilized on a CM5 chip with 7,600 RU and the wild-type (H3) and mutant histone H3 (H3K14A) peptides were injected at different concentrations (0, 7, 15, 32, 65, 125, 250 and 500nM). **(i)** The relative binding of H3 and H3K14A peptides toward Tt_Importin4 is shown in a bar graph. The average of the rations between the binding of the wild type peptide to that of the mutant peptide at each concentration are drawn in Y axis. The errors among the ratios were combined to generate the error bar.

Removing the carboxylate from Tt_E994 (Hs_E997) abolished the binding of the histone H3 tail, a result consistent with the crystal structure of the C-Importin4_histone H3 peptide complex. Although another carboxylate from Hs_D994 (Tt_D991) holds histone H3 K14, we were not able to examine the contribution of this residue to the binding due to insolubility of this mutant. We then mutated Tt_E881 (Hs_E884) and Tt_E834 (Hs_E837) to alanines and examined their binding (Fig. 4d, 4e, 4f). These mutants also showed defects in histone H3 binding, thus suggesting that the histone binding pocket in Importin4 observed in the crystal structure is indeed important in histone binding. We were able to observe four amino acids (_12_GGKA_15_) out of 18 amino acids

(_1_ARTKQTARKSTGGKAPRK_18_) used for crystallization, thereby indicating that histone H3 Lys14 might be a major residue interacting with Importin4. Therefore, we examined the contribution of histone H3 Lys14 to the binding with Importin4. We immobilized full-length Tt_Importin4 on a CM5 chip and used a wild-type (H3 Lys14, 1-27 a.a.) and a mutant (H3 Lys14Ala, 1-27 a.a.) histone H3 peptide as analytes (Fig. 4g, 4h and 4i). We measured the relative binding of the mutant peptides compared with the H3 Lys14 wild-type peptide. Mutating Lys14 to alanine significantly decreased, but did not completely abolish the binding to Importin4, which is consistent with the previous work by Soniat et al. [23]. These data suggest that histone H3 Lys14 plays a major role in the binding process and that other regions of the histone H3 peptide beside Lys14 also contribute to the binding of Importin4.

### Molecular architecture of the Importin4_Histone H3/H4_Asf1a complex through an integrative approach

One of the key questions in the DNA replication dependent chromatin assembly is to understand the mechanism of Importin4_histone H3/H4_Asf1 complex. To get closer to correct answer, we approach to visualize the 3D structure of whole complex using negative stained EM, SAXS and cross-linking mass spectrometry. Specifically, our XL-MS data on the inter-molecular interactions among full-length Importin4, histone H3, histone H4, or Asf1a, as well as the intra-molecular interactions within each protein provided the spatial proximity of the four protein components and their conformations (Fig. 1b, and Supplementary Table 1). In addition, the atomic structures of the C-Importin4 (668 to1,081 a.a.) from this study, as well as the structure of the histone H3/H4_Asf1a complex from other studies are available [7]. On the basis of these data, we computed the structural models of the full-length Importin4 bound to the histone H3/H4_Asf1a complex through an integrative modeling approach as implemented in the Integrative Modeling Platform (IMP) program [29] (Fig. 5a), combining all the data described above (see Methods for more details). The models of the complex relies on atomic structures determined by X-ray crystallography (C-Importin4_668-1081_ in this study and PDB ID: 2IO5 for histone H3/H4_Asf1a) [7], as well as comparative models of full-length Importin4 built with MODELLER 9.13 [30] (Material and Methods) based on the known structures (PDB ID: 3W3T) [28] detected by HHPred [31, 32] (Supplementary Fig. 6). While avoiding steric clashes and retaining sequence connectivity, 150,000 models of the complex were computed using Replica Exchange Gibbs sampling based on the Metropolis Monte Carlo algorithm. We only considered for further analysis an ensemble of the 500 best-scoring models (i.e., the ensemble), because there was effectively a single cluster of the 500 models, at the model precision of ~11 Å (Supplementary Fig. 7b and Materials and Methods). The resulting ensemble of the 500 models was validated on the basis of several considerations, as follows. First, the ensemble of best-scoring models satisfies 64 of the 72 cross-links within the Cα distance threshold of 35 Å (Fig. 2 and 4c; Supplementary Fig. 8), thus suggesting that the integrative model of the complex is consistent with the XL-MS data as well as X-ray crystallographic structures. Second, all models in the ensemble satisfy the excluded volume and sequence connectivity restraints used to compute the models. Third, we performed negative-stain single-particle EM analysis on the Importin4_histone H3/H4_Asf1a complex. We collected a total of 12,725 particles from EM images, and further processed the data to generate 2D class-averages. We then compared each image of the class-averages with the best-matching projection computed from the models in the ensemble. The models of the complex also fit the EM class averages, revealing that our complex model is consistent with the overall shape of the complex obtained by the negative-stain EM analysis (Fig. 5b). Finally, we collected a SAXS profile of the Importin4_histone H3/H4_Asf1a complex (Supplementary Fig. 9). The molecular weight of the SAXS sample was estimated to be 162.1 kDa by using SAXS MOW [33], a value nearly identical to the expected molecular weight of 161 kDa from the sequence (thus confirming the monomeric state of the SAXS sample). Additionally, the linearity in the Guinier plot confirms a high degree of compositional homogeneity of the complex in solution (Supplementary Fig. 9a). The radius of gyration (*R_g_*) and the maximum particle size (*D_max_*) were determined as 41.9 ± 0.2 and 140.5 Å, respectively (Supplementary Fig.9a,c). Subsequently, the model of the complex shows that the entire HEAT repeats of Importin4 wrap histone H3/H4_Asf1a with extensive interactions with histone H3/H4. In agreement with this model, the C-terminal region of Importin4 alone cannot form a stable complex with histone H3/H4_Asf1a (unpublished data). Thus, we present the molecular architecture of the Importin4_Histone H3/H4_Asf1a complex produced through the integrative modeling approach, based on data from the X-ray crystallography, XL-MS, EM, and SAXS.

**Fig. 5.**
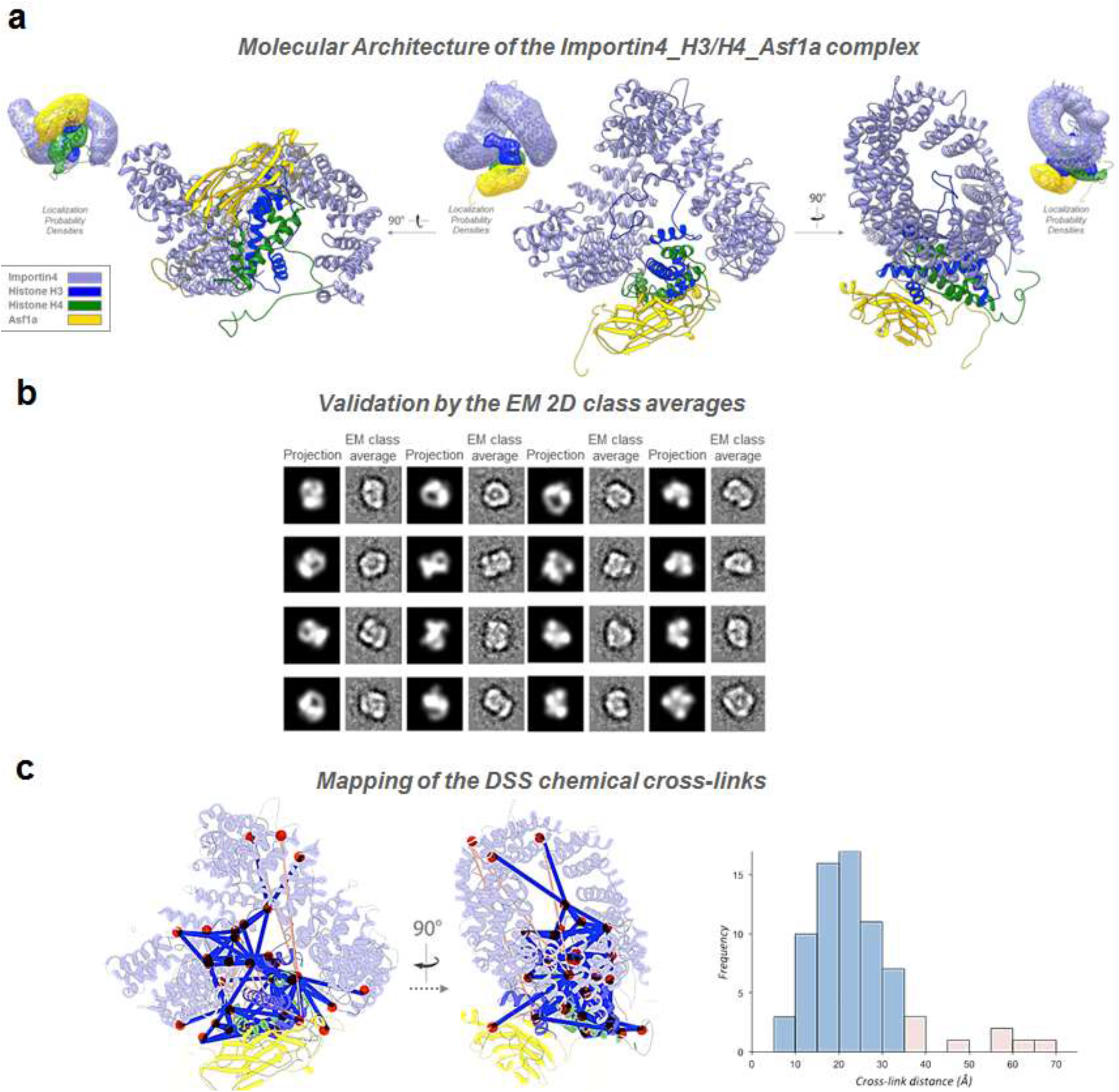
The molecular architecture of the Importin4_Histone H3/H4_Asf1a complex produced through an integrative modeling approach. **(a)** Three views of the representative model (the centroid in the ensemble) of the Importin4_histone H3/H4_Asf1a complex are shown in ribbon representation. In all views, the components of the complex are colored in pale purple (Importin4), blue (histone H3), green (histone H4), and yellow (Asf1a). At a side, the localization probability density map is shown along with the representative model in it; the ensemble of the 500 best-scoring models was converted into the localization probability density map that specified how often grid points of the map were occupied by a given protein. **(b)** Correspondence of the negative-stain EM 2D class averages and the model projections. Each of the EM class averages is shown, along with its best-matching projection computed from the ensemble of the complex models. For each diagram, the projection is shown on the left side, and the corresponding EM class average is shown on the right side. **(c)** The ensemble of the complex models match well with the chemical cross-linking data. The chemical cross-links falling within the C_α_-C_α_ distance threshold of 35 Å (blue lines) or outside of that threshold (orange lines) are shown on the representative model of the complex (ribbon representation). Each of the cross-linked lysine residues is depicted as a red circle. The histogram shows the C_α_-C_α_ distance distribution of all 72 DSS cross-links in the model, suggesting that the integrative model of the complex is consistent with the XL-MS data as well as X-ray crystallographic structures.

## Discussion

Here, we provide structural insights into how Importin4 recognizes the histone H3 tail and the molecular architecture of the entire Importin4_histone H3/H4_Asf1a complex. During DNA duplication, many histones need to be synthesized, transported into the nucleus, and eventually assembled into nucleosomes. Histones are highly charged proteins and contain strongly-hydrophobic surfaces. Because of these properties, histones can bind to other proteins nonspecifically. Therefore, newly synthesized histones are escorted by several histone chaperones in the cytoplasm as well as the nucleus. Importin4 receives histone H3/H4 from the cytosol, transfers the histone into the nucleus, and eventually nuclear histone chaperones accept the histone H3/H4 for nucleosome [34–38]. During this nuclear import, the hydrophobic region of the histone H3/H4 dimer remains protected by the histone chaperone Asf1a [7, 8]. Our XL-MS data show that Importin4 interacts with the DNA binding surface of histone H3/H4 which is masked by other histone chaperones in the cytosol. Therefore, it appears that Importin4 not only recognizes the NLS of the histone H3 for nuclear transport but also protects the highly-charged DNA binding surface with Asf1 during nuclear import. Notably, we have observed cross-links between Importin4 and the histone fold of H3/H4, and between Importin4 and Asf1a. These data suggest that Asf1a and the histone fold as well as histone tail might contribute to the binding to Importin4 together with histone H3/H4. Histone H3 and H4 form a dimer when they are transported into the nucleus [4, 7, 8, 39]. The NLS of histone H3 is located at the N-terminal tail and histone H3 Lys14 is shown to be important for the function of the histone H3 NLS in yeast [19]. Furthermore, recent studies have shown that histone H3 Lys14 is a major binding residue for several other importins including human Importin4 [23, 24]. Consistent with these previous data, our XL-MS data showed that Importin4 binds extensively to the N-terminal tail of histone H3. From our crystal structure, we were only able to resolve a four amino acid segment (_12_GGKA_15_) including the histone H3 Lys14 although we used the 18 a.a. histone H3 tail (_1_ARTKQTARKSTGGKAPRK_18_). This observation further supports that histone H3 Lys14 is a major binding residue in the histone H3 tail. The N-terminal tail of histone H4 also serves as a NLS signal in yeast [10, 19]. Therefore, the NLSs of histone H3 and H4 appear to be functionally redundant. However, it is unclear whether the same Importin is responsible for recognizing the NLSs of histone H3 and H4. Our limited proteolysis analysis showed that the N-terminal tail of histone H3 but not histone H4 is protected in presence of Importin4. Therefore, it seems that Importin4 predominantly recognizes the histone H3 tail as a NLS. Although our XL-MS data and the crystal structures shows that the C-terminal region of Importin4 is a major domain responsible for interacting with the histone H3/H4_Asf1a complex, it is unclear whether the N-terminal region of Importin4 is also involved in recognizing the NLS of histones or interacting with histone H3/H4_Asf1a. Because C-Importin4 was not able to form a stable complex with histone H3/H4_Asf1a in our hands (unpublished data), it is likely that the N-terminal region of Importin4 is also involved in recognizing the histone H3/H4_Asf1a complex.

For the binding analysis, we used dolphin Importin4 (Tt_Importin4), because we were not able to obtain a stable full-length human Importin4 in the absence of histone H3/H4_Asf1a. Although humans and dolphins are evolutionarily distant, the sequence conservation between Importins is remarkably high, with 89% identity. Interestingly, when we compared the affinities of C-Importin4 and full-length Tt_Importin4, we found that the full-length Tt_Importin4 shows much slower Kon and Koff for histone H3 binding. These data suggest that the N-terminal region of Importin4 might be involved in recognizing the histone H3 NLS as well as the histone H3/H4-Asf1 complex. It is unlikely that the different binding between full-length Tt_Importin4 and C-Importin4 could result from the species difference, because the sequence identity between the two is so high. However, it is unclear which region (if any) in the N-terminal of Importin4 might have a second binding site for histone H3.

In this study, we obtained four different types of experimental data for the Importin4_histone H3/H4_Asf1a complex, as described above; (1) 72 intra- and inter-molecular DSS cross-links were identified via mass spectrometry (Fig. 1), identifying the spatial proximities among the four protein components and their conformations (Fig. 4c); (2) an atomic structure of the Importin4 C-terminal domain (668 to1081 a.a.) (Fig. 2) was determined by X-ray crystallography, informing its representation; [40] 2D class average images of the Importin4_histone H3/H4_Asf1a complex were obtained by negative-stain EM, delineating the shape of the complex (Fig. 4b); (4) a SAXS profile was collected, outlining the shape and potential dynamics of the complex (Fig. 4d). By combining all of these data, we computed and validated the ensemble of structural models of the Importin4_histone H3/H4_Asf1a complex that satisfy all information. The EM and SAXS analysis showed that the resulting ensemble of the complex models reflects the molecular architecture of the complex *in vitro*, suggesting the potential interactions among Importin4, histones, and Asf1a.

Importin4 is composed of HEAT repeats conferring conformational flexibility. Previously, X-ray crystallography and SAXS experiments showed that Importin-β, another HEAT repeat protein, undergoes conformational change upon binding to its cargoes and binding partners [41]. Consistent with this, a plateau in the Kratky plot (q=0.15 - 0.25 Å^-1^, Supplementary Fig. 9b) suggests some flexibility in the complex and we were also able to observe an extended conformation of the complex in the 2D class averages obtained from negative-stain EM (Supplementary Fig. 10).

In summary, our study delineates the interactions among Importin4, histone H3 and Asf1a providing mechanistic insights into how Importin4 recognizes the histone H3 NLS. In addition we provide the molecular architecture of the Importin4_Histone H3/H4_Asf1a complex produced through the integrative modeling approach that combines X-ray crystallography, XL-MS, SAXS, and EM.

## Materials and methods

### Protein Expression and Purification

A gene encoding the C-terminal region of human Importin4 (C-Importin4) (residues 668 to 1081 a.a.) was cloned into a modified pET28a vector with a TEV cleavage site after a His-tag. Cells expressing C-Importin4 were grown to an optical density of 0.8 at 600 nm, induced with 0.5 mM Isopropylthiogalactoside and grown at 18 °C for 16 hrs before being harvested. The cells were resuspended in a buffer A (50mM Tris-HCl pH 8.0, 500mM NaCl, 5% glycerol) and lysed by sonication. The cell lysate was centrifuged at 18,000 rpm for 40 min. The supernatant was then incubated with Ni-NTA resin (Qiagen). The resin was washed in a buffer B (50mM Tris-HCl pH 8.0, 500mM NaCl, 5% glycerol and 20mM imidazole) and eluted in a buffer C (50mM Tris-HCl pH 8.0, 500mM NaCl, 5% glycerol and 100mM Imidazole). The eluted protein was dialyzed overnight at 4 °C in the presence of TEV protease in a buffer D (50mM Tris-HCl pH 8.0, 100mM NaCl, 1mM EDTA, 1mM DTT) to remove the N-terminal His tag. Uncleaved proteins were recaptured on the Ni-NTA resin, and the proteins were further purified by a Q column (GE Healthcare Life Sciences) by salt-gradient elution from 50mM NaCl to 1M NaCl, and by Superdex 200-26-60 (GE Healthcare Life Sciences) size-exclusion chromatography in a buffer E (50mM Tris-HCl pH 8.0, 500mM NaCl, 5mM DTT). Tt_Importin4 and mutant proteins were purified similarly and stored in a buffer (20mM HEPES pH 7.5, 100mM NaCl and 2mM TCEP). Se-Met labeled proteins were also generated in the same way as the wild-type proteins.

### Cross-Linking Mass Spectrometry

The Importin4_H3/H4_Asf1a complex (50 g, approx. 1 mg/ml) was mixed with 25 mM disuccinimidyl suberate [25]-d0/d12 (Creative Molecules) to a final concentration of 1 mM. The mixture was incubated for 30 min at 37 °C with mild shaking (700 rpm) on a thermomixer (Eppendorf). The crosslinking reaction was quenched by adding ammonium bicarbonate (Sigma-Aldrich) to a concentration of 50 mM. The cross-linked samples were denatured with 8M urea (Sigma-Aldrich) in the presence of 2.5 mM tris(2-carboxyethyl) phosphine. The free cysteine thiols were blocked by adding iodoacetamide (Sigma-Aldrich) to 5 mM final concentration. The samples were then sequentially digested with Lys-C (3 hrs) and trypsin (overnight) proteases at 1:100 and 1:50 enzyme-to-substrate ratio, respectively. The cross-linked peptides were enriched by SEC (Superdex Peptide 3.2/30, GE Healthcare Life Sciences) as described previously [42] and analyzed on an Orbitrap Elite mass spectrometer (Thermo) XL mass spectrometer (Thermo). Data analysis was performed using xQuest [43], and a sequence database containing the four target proteins. A score cut-off of 16 was used and all spectral exceeding this threshold were manually evaluated. The estimated false discovery rate of the cross-links is 5%, as determined by xProphet (Walzthoeni et al., 2012).

### Limited Proteolysis

Proteins (100 μg) in 50 mM Tris-HCl (pH 8.0) containing 100 mM NaCl were incubated with trypsin (2 μg) at 37 °C for 60 mins. The reaction was terminated at several different time points by the addition of SDS-PAGE sample loading buffer. The proteins were resolved on a 15% SDS polyacrylamide gel and visualized by staining in Coomassie brilliant blue. The bands were identified with N-terminal sequencing.

### Structure Determination of the C-Importin4 Alone and the C-Importin4 Bound to the NLS of the Histone H3

The crystals of C-Importin4 (residues 668 to 1081 a.a.) were grown with a hanging drop vapor diffusion method at 18 °C, by mixing 1μl of the protein with 1 μl reservoir solution containing 240mM sodium tartrate dibasic dehydrate and 20% PEG 3350. For the complex crystals, histone H3 peptide (residues 1 a.a to 18 a.a) (Anygen) and C-Importin4 (residues 668 to 1081 a.a) were mixed at a molar ratio of 1:5 and grown with a hanging drop vapor diffusion method at 18 °C, and the crystals were obtained in the same condition as for C-Importin4 alone. The crystals were soaked in a cryoprotectant with additional 20% glycerol (v/v) added to the mother liquor condition, and frozen in liquid nitrogen. X-ray diffraction data were collected at the NW12A beamline in the Photon Factory, Tsukuba Japan. The data were indexed, integrated, and scaled using HKL2000 [44]. The structures of C-Importin4 alone and the peptide complex were determined by SAD using an Se anomalous signal, and molecular replacement with a search model of C-Importin4 respectively, using Phaser [45]. Refinement was performed using CNS [46] and Phaser, and the models were built with Coot [47] or O programs. The structure factors of C-Importin4 and the C-Importin4_histone H3 peptide complex were deposited in the PDB database with PDB ID 5XAH, and 5XBK respectively.

### Affinity Measurement Using Surface Plasmon Resonance

Surface Plasmon Resonance (SPR) experiments were performed using a Biacore T200 instrument (GE Healthcare Life Sciences) at 7 °C. C-terminal biotinylated histone H3 peptides (1-35 a.a.) were synthesized at Anygen, Korea, and coupled on a CM5 chip coated with streptavidin. The wild-type and mutant proteins were injected at a flow rate of 30 l/min using the binding buffer (10 mM HEPES, 150 mM NaCl) and the binding was measured at various concentrations at 7 °C. The CM5 chip was regenerated at a flow rate of 30 l/min using a regeneration buffer (1M NaCl and 20 mM NaOH in binding buffer) after each injection. For the binding analysis with wild-type and mutant histone peptides, Tt_Importin4 was immobilized on a CM5 chip and the histone H3 peptides were injected at a flow rate of 30 μl/min. The binding was measured at various concentrations at 7 °C. The chip was regenerated at a flow rate of 30 μl/min using the regeneration buffer (1M NaCl and 20 mM NaOH in binding buffer). To compare the relative bindings, RU values at four different concentrations (125, 250, 500, 1000 nM) of the wild-type and the mutants were collected. The ratios between the RU values of the mutants and the wild-type at each concentration were calculated. The error bars were generated from the four ratios calculated from four different concentrations for each mutant. For the histone H3 deletion peptides, the peptide of H3_1-18_: 1 to 18 a.a, H3_1-11_: 1 to 11 a.a. and H3_1-6_: 1 to 6 a.a. were synthesized at Anygen, Korea. C-Importin4 was immobilized on a CM5 chip and the histone H3 deletion peptides were injected at a flow rate of 30 μl/min. The binding was measured at various concentrations (0, 0.5, 1, 2, 4, 6 and 8 uM) at 7 °C. The chip was regenerated at a flow rate of 30 μl/min using the regeneration buffer (500 mM NaCl in binding buffer).

### Negative-stain Electron Microscopy

The purified Importin4_H3/H4_Asf1a complex was applied to ultracentrifugation at 74,329 xg for 16 hr with a 5-20% sucrose and 0-0.2% glutaraldehyde gradient. The fractionated complex was negatively-stained with 2% (w/v) uranyl acetate for 1 minute on 400 mesh carbon-coated Cu grids. Images were recorded at 50,000x magnification with a defocus value of 0.8 - 2.3 μm on a 4x4K TVIPS (Tietz Video and Imaging Processing System) CCD camera, attached to a Jeol 2100F field emission gun transmission electron microscope at 200 kV. The data set was further processed using the EMAN2 program software package [48]. A total of 13,398 particles were boxed by e2box.py from 52 micrographs and then subjected to reference-free 2D classification. The ensemble of the complex models was projected to 210 images by e2proc3d.py to compare with the EM 2D class averages. The similarity comparison between the set of 2D class averages and projections was computed by e2simmx.py. Then, best matching values were searched from the similarity matrix and presented by e2classify.py. The best 16 matches were selected, and projections were then aligned to their partner classes in the same direction by e2align2d.py.

### Integrative Structure Modeling of the Importin4_H3/H4_Asf1 Complex

Integrative structure modeling of the Importin4_H3/H4_Asf1a complex proceeded through four stages [29, 49–51]: (1) gathering data, (2) representing subunits and translating the data into spatial restraints, [40] configurational sampling to produce an ensemble of structural models that satisfies the restraints, and (4) analyzing and validating the ensemble models and data. The modeling protocol (*i.e*., stages 2, 3, and 4) was scripted using the *Python Modeling Interface* (PMI) package, version 4d97507, a library for modeling macromolecular complexes on the basis of the open-source *Integrative Modeling Platform* (IMP) package, version 2.6 (http://integrativemodeling.org) [29].

#### Stage 1: Gathering data

During this study, we obtained four different types of experimental data for the Importin4_H3/H4_Asf1a complex, as described above; (1) 72 intra- and inter-molecular DSS crosslinks were identified *via* mass spectrometry (Fig. 1), identifying the spatial proximities among the 4 protein components and their conformations (Fig. 4c); (2) an atomic structure of the Importin_4668-1081_ CTD domain (Fig. 2) was determined by X-ray crystallography, informing its representation; [40] 2D class average images of the Importin4_H3/H4_Asf1a complex were obtained by negative-stain EM, delineating the shape of the complex (Fig. 4b); and (4) a SAXS profile was collected, outlining the shape and potential dynamics of the complex (Fig. 4d).

#### Stage 2: Representing subunits and translating data into spatial restraints

Representations of the complex relied on atomic structures determined by X-ray crystallography (Importin_4668-1081_ in this study and PDB ID: 2IO5) [7] as well as comparative models built with MODELLER 9.13 [30] based on the known structures detected by HHPred [31, 32]; HHPred detected a template structure of *Saccharomyces cerevisiae* karyopherin Kap121p (PDB ID: 3W3T) [28] as a-92 best-matching template structure (Probability=100.00, E-value=4.2x10^−92^, Score=910.68, Sequence Identities=19%, and Sequence Similarity=0.243; Supplementary Fig. 6). We then used the template structure for comparative modeling of Importin4^1-667^, using MODELLER 9.13 [30]; 100 models were created and a structural model that minimized the DOPE score (DOPE=-125084.10938) [52] was selected subsequently for the integrative modeling of the Importin4_H3/H4_Asf1a complex.

The components of the Importin4_H3/H4_Asf1a complex were coarse-grained using beads representing either a rigid body or a flexible string, based on the available X-ray structures and comparative models, as follows.

(1) *Rigid bodies*: A rigid body was defined for each X-ray structure and comparative model. Rigid bodies were coarse-grained using two resolutions, where beads represented either individual residues or segments of 10 residues. For the one-residue bead representation, the coordinates of a bead were those of the corresponding C_α_ atoms. For the 10-residue bead representation, the coordinates of a bead were the center of mass of all atoms in the corresponding consecutive residues. In a rigid body, the beads have their relative distances constrained during configurational sampling [50]. It has been known that the N-terminal and C-terminal regions in the importin family undergo conformational changes relative to each other in various conditions [53, 54]. Therefore, we represented each of the N-terminal (residues 1-531) and the C-terminal (residues 532-1081) regions of Importin4 as a separate rigid body to allow their relative motions during sampling, based on normal mode analysis using HingeProt [55]. It is also supported by our investigation that the N-terminal and C-terminal regions hold nearly identical conformations of helices on their own, despite of apparent variability in their relative orientations among PDB structures of the related importin family.
(2) *Flexible strings of beads*: The remaining regions without an atomic model (*i.e.*, the predicted disordered regions) were coarse-grained by flexible strings of beads representing up to 10 residues. In a flexible string, the beads are restrained by the sequence connectivity and excluded volume restraints [50].

After obtaining this representation, we next encoded the spatial restraints into a scoring function based on the information gathered in Stage 1, as follows.

(1) *Cross-link restraints:* The 72 DSS cross-links were used to restrain the distances spanned by the cross-linked residues [50, 56]. The cross-link restraints were applied to the corresponding residue pairs. Each cross-link restraint functions effectively as an upper-bound harmonic restraint of ~17 Å on the distance between the cross-linked residues.
(2) *Excluded volume restraints*: The protein excluded volume restraints were applied to each 10-residue bead, using the statistical relationship between the volume and the number of residues that it covered [49, 52, 56].

[40] *Sequence connectivity restraints*: We applied sequence connectivity restraints, using a harmonic upper distance bound on the distance between consecutive beads in a subunit, with a threshold distance equal to the sum of the radii of the two connected beads. The bead radius was calculated from the excluded volume of the corresponding bead, assuming standard protein density [49, 50, 56].

#### Stage 3: Configurational sampling to produce an ensemble of structural models satisfying the restraints

The search for good-scoring models relied on Replica Exchange Gibbs sampling, based on the Metropolis Monte Carlo algorithm [50, 51]. The Monte Carlo moves included random translation and rotation of rigid bodies (up to 2 Å and 0.04 radians, respectively), and random translation of individual beads in the flexible segments (up to 3 Å). Up to 8 replicas were used, with temperatures ranging between 1.0 and 2.5. Twenty independent sampling calculations were performed, each one starting with a random initial configuration. A model was saved every 10 Gibbs sampling steps, each consisting of a cycle of Monte Carlo steps that moved every rigid body and flexible bead once. The sampling produced a total of 100,000 models in 20 independent runs. We only considered for further analysis an ensemble of 500 best-scoring models with a score better than 295 (this threshold ensured satisfaction of the restraints used to compute the models; see below).

#### Stage 4: Analysis and validation of the ensemble models and data

Input information and output models need to be analyzed to estimate model precision and accuracy, detect inconsistent and missing information, and to suggest more informative future experiments. We used the analysis and validation protocol published earlier [49, 51]: Assessment began with a test of the thoroughness (convergence) of sampling, followed by clustering and variability in the ensemble of best-scoring models, quantification of the model fit to the input information, and assessment by data not used to compute the structural model.

##### (1) Convergence of sampling

First, we assessed the thoroughness of the configurational sampling by comparing a subset of 250 best-scoring models from runs 1–10 to another subset of 250 best-scoring models from runs 11–20. Each subset of the best-scoring models was converted into a density map that specified how often grid points of the map were occupied by a given protein (the “localization probability density map”). Importantly, the two localization probability density maps were nearly identical to each other (crosscorrelation coefficient = 0.99; Supplementary Fig. 7a), demonstrating that the Monte Carlo algorithm well sampled all models that satisfied the input restraints. Consistently, the TM-score [57], a measure of structural similarity, between representative (centroid) models of the two subsets is 0.951, indicating about the same fold.

##### (2) Clustering and variability in the ensemble of best-scoring models

K-means clustering [50] identified a single dominant cluster containing 397 similar models (79.4%) from the ensemble of 500 best-scoring models (Supplementary Fig. 7b, c). The largest root-mean-square deviation (RMSD) between a pair of the 500 models was ~15 Å. However, the most of the RMSD values were less than ~11 Å (Supplementary Fig. 7b); more specifically, the most of the RMSD values between the dominant (397 models, 79.4%) and minor clusters (103 models, 20.6%) were relatively low at less than ~11 Å, considering uncertainty of the data and the coarse-grained representation of some unstructured regions. The TM-score [57] between representative (centroid) models of the two clusters is 0.889, indicating about the same fold. The cross-correlation coefficient between the localization probability densities of the two clusters is 0.99 (Supplementary Fig. 7c). As a result, the localization of all protein components is nearly identical between the dominant and minor clusters. Therefore, we conclude that there is effectively a single cluster of the 500 best-scoring models, at the model precision of ~11 Å (equal to the RMSD threshold above).

For the remainder of our analysis, we used all of the 500 structural models. The localization probability density maps were computed from the ensemble of the 500 best-scoring models (Fig. 4a). The positional variability in the placement of H3/H4/Asf1b relative to Importin4 is relatively small within ~10 Å, consistent with the localization probability density maps (Fig. 4a). The representative model in Fig. 4 and Supplementary Fig. 7–8 was chosen from the centroid model in the ensemble.

##### [40] Fit to input information

An accurate structural model needs to satisfy the input information used to compute it. The ensemble of models was assessed in terms of how well the models satisfied the information from which they were computed, including the chemical cross-links, excluded volume, and sequence connectivity restraints, as follows.

The ensemble of the 500 best-scoring models satisfied 89% of the DSS cross-links (Fig. 4c and Supplementary Table 1); a cross-link restraint was satisfied by the ensemble if the corresponding C_α_-C_α_ distance in any model in the ensemble was less than 35 Å. The 35 Å distance threshold was used only for the assessment of the cross-links. The chosen cut-off for classifying the cross-links as compatible with the model was selected with a bin size of 5 Å, taking into account the expected accuracy of the model. The histogram (Fig. 4c) presents nearly ideal Gaussian distribution for the cross-links, supporting our cross-link restraints worked well during sampling. Most of the cross-link violations were small, and could be rationalized by local structural fluctuations and coarse-grained representations of some unstructured regions (Supplementary Fig. 8). In addition, there are some general limitations of the XL-MS method that can lead to 5~10% of the cross-links violating a structure or a model. This is due to two main reasons: First, interpretation of the mass spectrometry data (matching theoretical with experimental data) is associated with an error rate that can be relatively easily controlled [42, 43], although this is not trivial for smaller databases. The approximate error rate (*i.e.* false discovery rate) for the data sets used in this work is 5%. Additional, non-compliant crosslinks may be observed in case of sample heterogeneity; the complex under study may be present in different conformational states or may be partially dissociated into subcomplexes. Because XL-MS is an ensemble method, the cross-links may therefore represent several such states, potentially leading to additional 5% violation of the cross-links. Therefore, the ensemble of the 500 best-scoring models satisfies the chemical cross-linking data within its uncertainty.

All best-scoring models in the ensemble satisfied the excluded volume and sequence connectivity restraints under the combined score threshold of 20.

##### (4) Satisfaction of data that were not used to compute structural models

The most direct test of a model is by comparing it with the data that were not used to compute the model (a generalization of cross-validation).

First, best-matching projections of the representative model (the centroid in the ensemble) fit the negative-stain EM 2D class averages (Fig. 4b). The class averages provide overall shape and size information of the complex when viewed from various angles. Our integrative model of the complex was validated by comparing each image of the EM 2D class averages with the best-matching projection that is computed from the model. As a result, the EM 2D analysis supports that the structural model is consistent with the overall shape and size of the complex obtained by the negative-stain EM analysis.

Second, our models in the ensemble are in agreement with the SAXS profile and *ab initio* shape (envelope) of the complex (Fig. 4d and Supplementary Fig. 9; see the Results section).

### Accession numbers

The structure factors for C-Importin4 and C-Importin4_Histone H3 peptide were deposited at PDB database (www.rcsb.org) as 5XAH and 5XBK respectively. The integrative model of the Importin4_H3/H4_Asf1a complex is available in PDB-Dev (https://pdb-dev.rcsb.rutgers.edu/).

## Acknowledgments

We thank all members of the Song Lab, T. Matsui and T.M. Weiss at the SSRL, SLAC National Accelerator Laboratory, for assistance in collecting SAXS data. This work was partially supported by grants (NRF-2016R1A2B3006293, NRF-2016K1A1A2912057, NRF-2014K2A3A1000137, 2011-0031955, CMCC) to J.S., grants (1-16-JDF-017, N019154-00, AG050903 and DK111465) to U.-S.C., and ERC grant Proteomics4D (AdvG grant #670821) to R.A. T.Y. is a recipient of The Jonasson Fellowship. This work was supported by The Jonasson Center for Medical Imaging and the Karolinska Institutet Center for Innovative Medicine.

## Abbreviations

NLS, Nuclear Localization Sequence; EM, Electromicroscopy; XL-MS, Crosslinking Massspectrometry; SAXS, Small Angle X-ray scattering; DSS, Disuccinimidyl suberate

---

**Supplementary Fig. 1.**
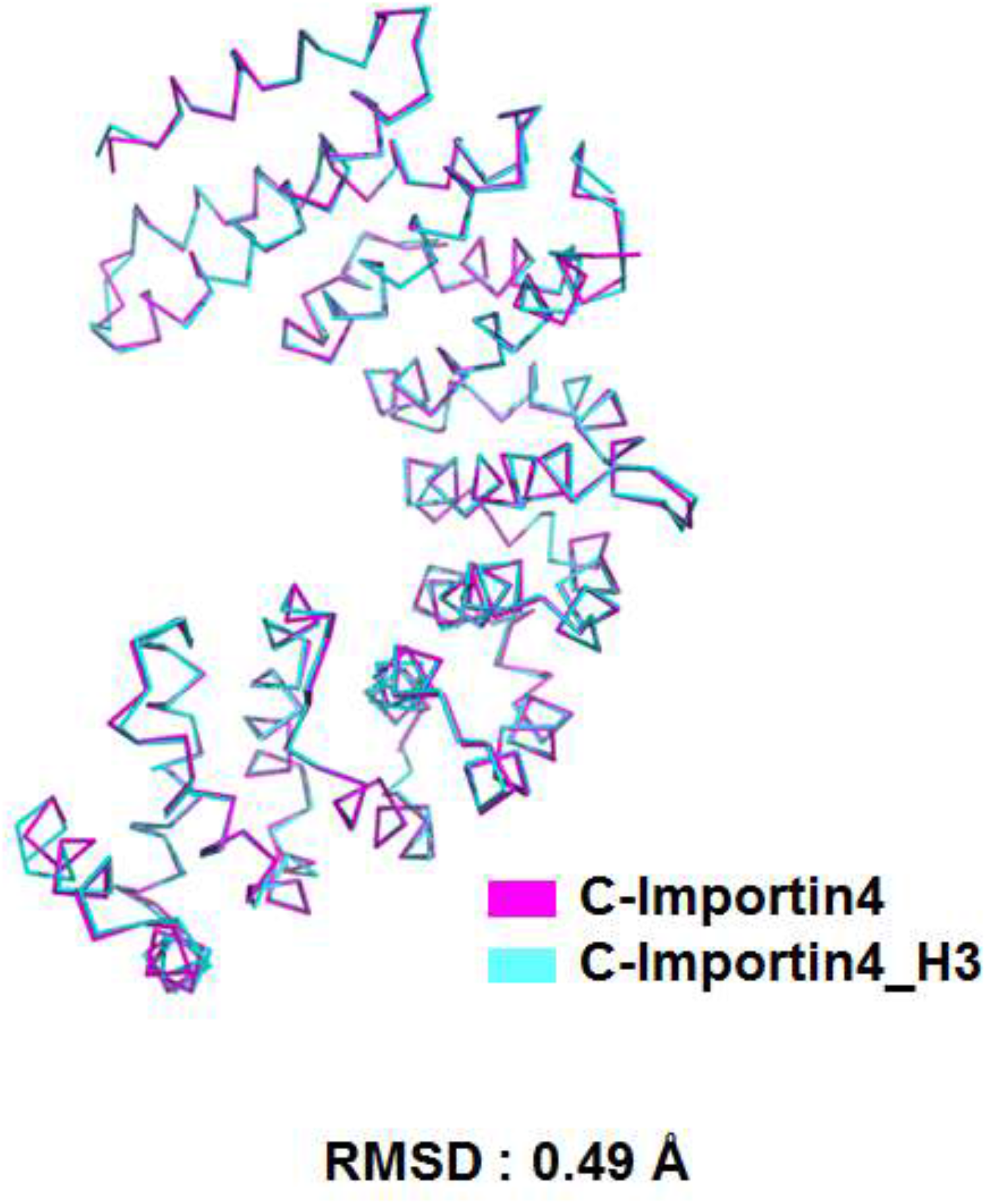
Superimposition of C-Importin4 and C-Importin4_H3 complex. C-Importin4 in magenta and C-Importin4_Histone H3 in cyan were superimposed with RMSD value is 0.49 Å.

**Supplementary Fig. 2.**
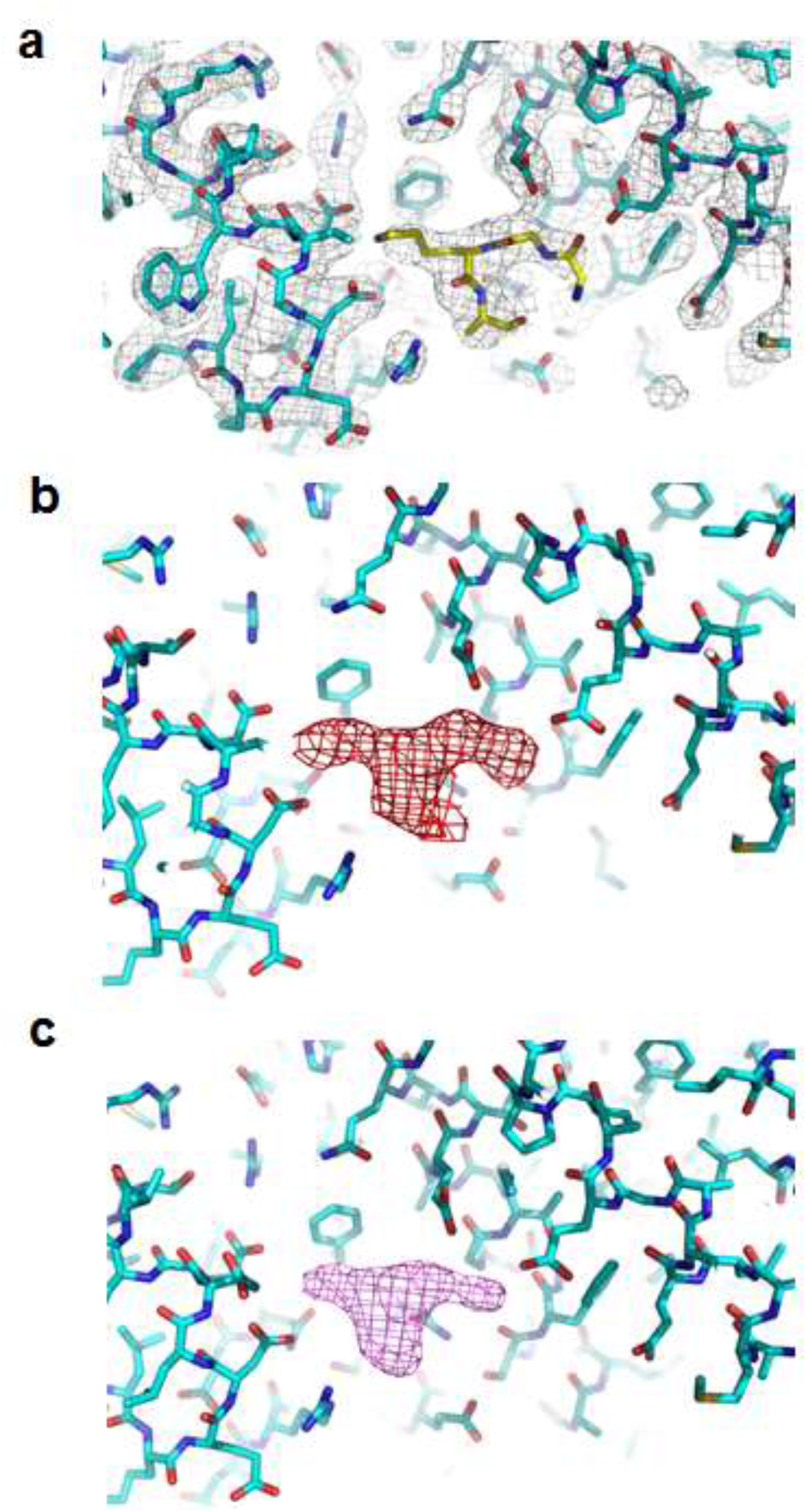
Electron density map near histone H3 peptide. (a) Electron density maps near the histone H3 peptide binding pocket. A 2F_o_-F_c_ map (1σ level) generated in presence of the peptide (b) a F_o_-F_c_ map (3 σ level) and (c) a kicked omit map (3 σ level) generated in the absence of the peptide.

**Supplementary Fig. 3.**
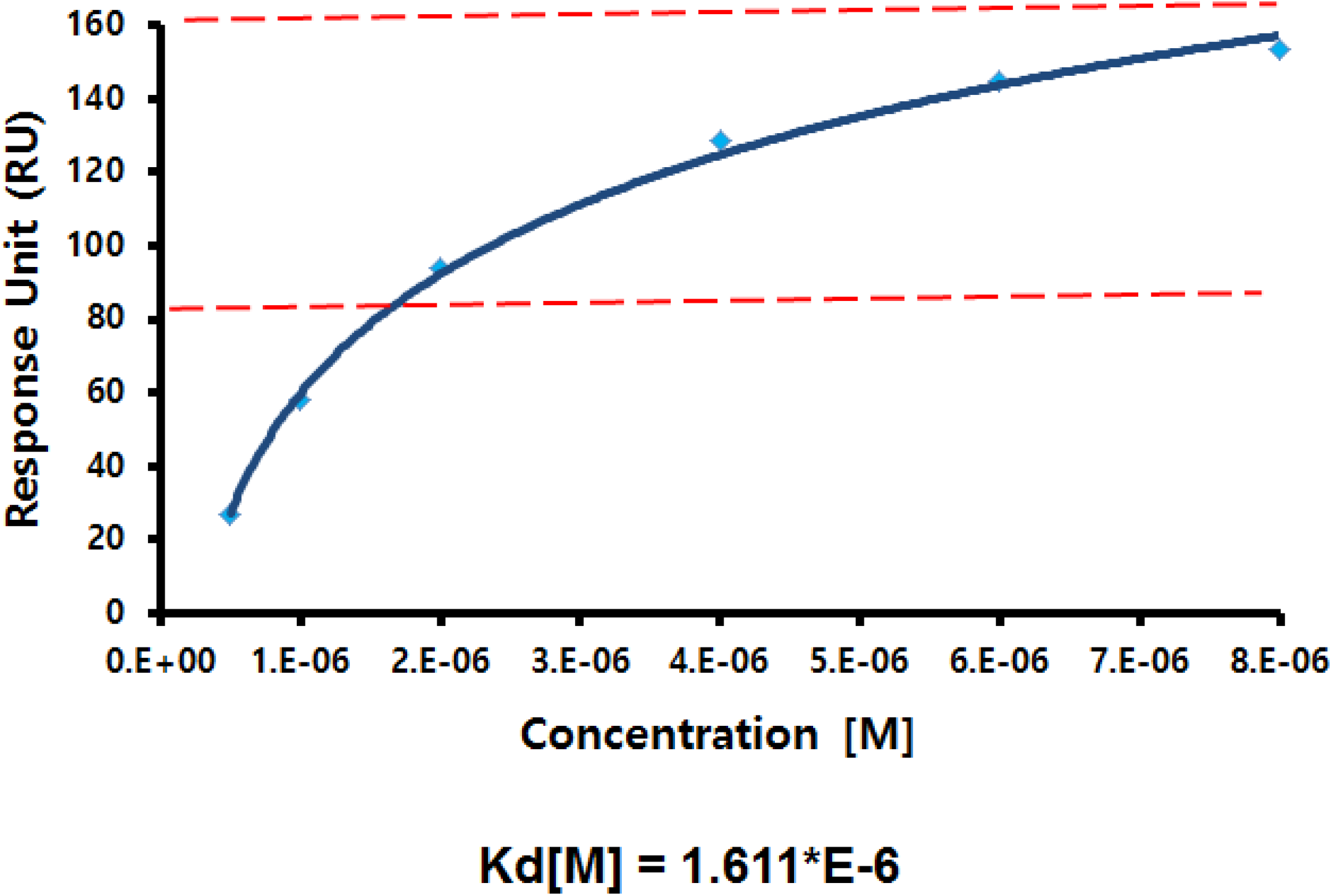
SPR analysis of C-Importin4 binding with histone H3 peptide. The K_D_ between C-Importin4 and histone H3 peptide was estimated by plotting the response units from Figure 3A (Y-axis) versus the concentrations (0.5, 1, 2, 4, 6 and 8 μM) of C-Importin4 (X-axis). The concentration at the half maximum of RU was estimated as 1.6 μM.

**Supplementary Fig. 4.**
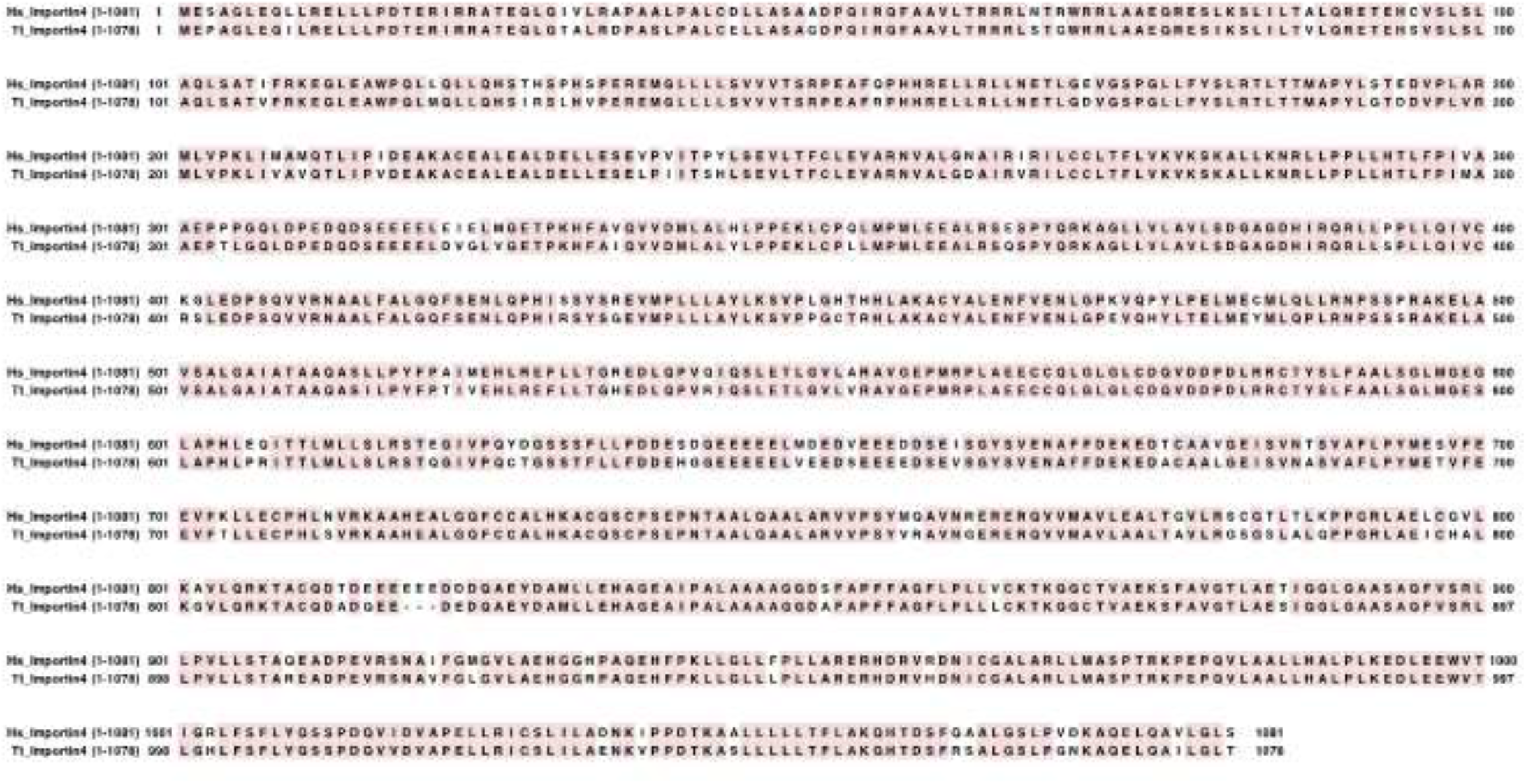
Sequence alignment between human Importin4 and dolphin Importin4. The upper lane indicates *Homo sapiens* (Hs) Importin4, and the lower lane indicates *Tursiops truncatus* (Tt) Importin4.

**Supplementary Fig. 5.**
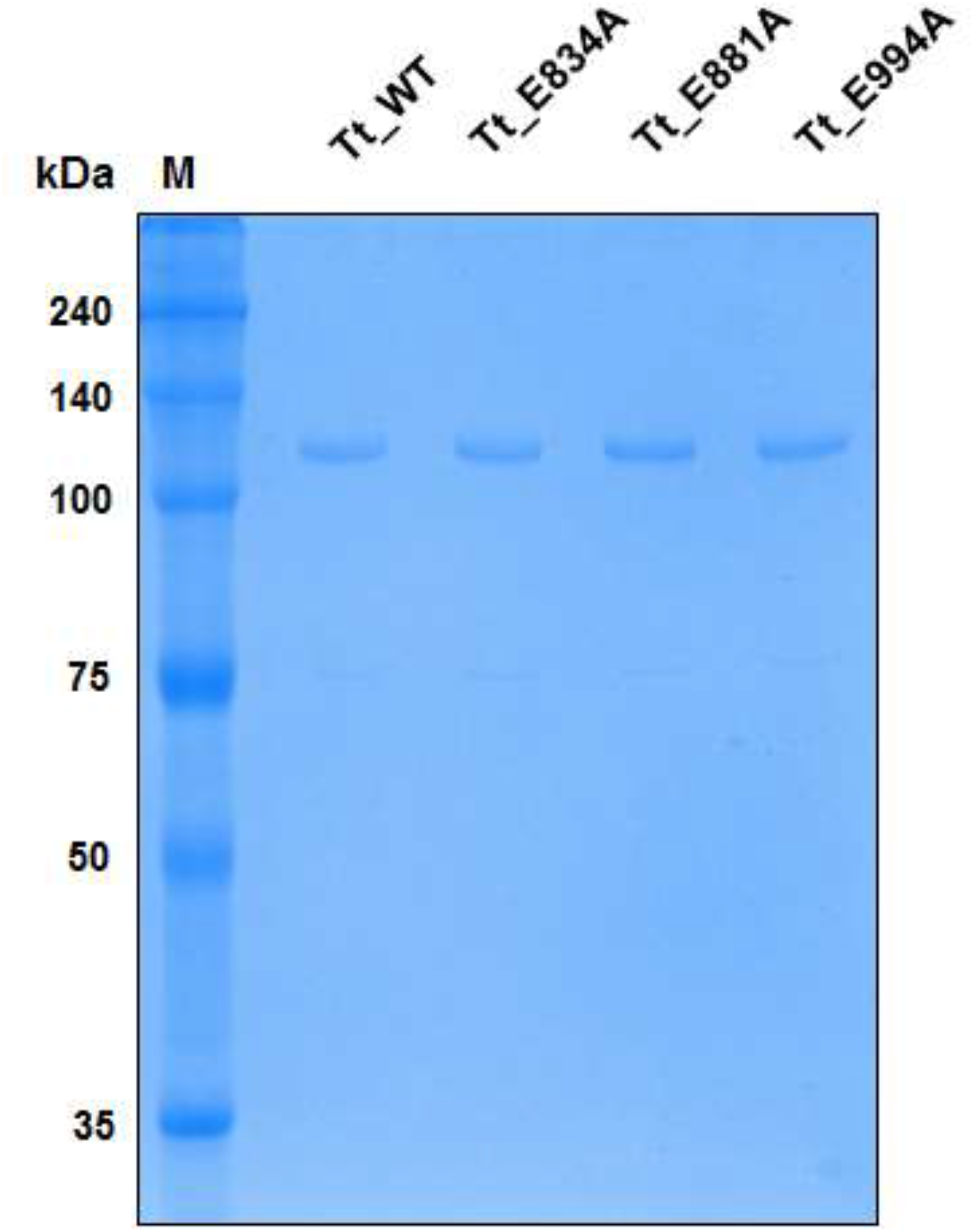
SDS-PAGE analysis of Tt_Importin4 proteins. A SDS-PAGE shows Wild-type Tt_Importin4 and three Tt_Importin4 mutants used for SPR analysis.

**Supplementary Fig. 6.**
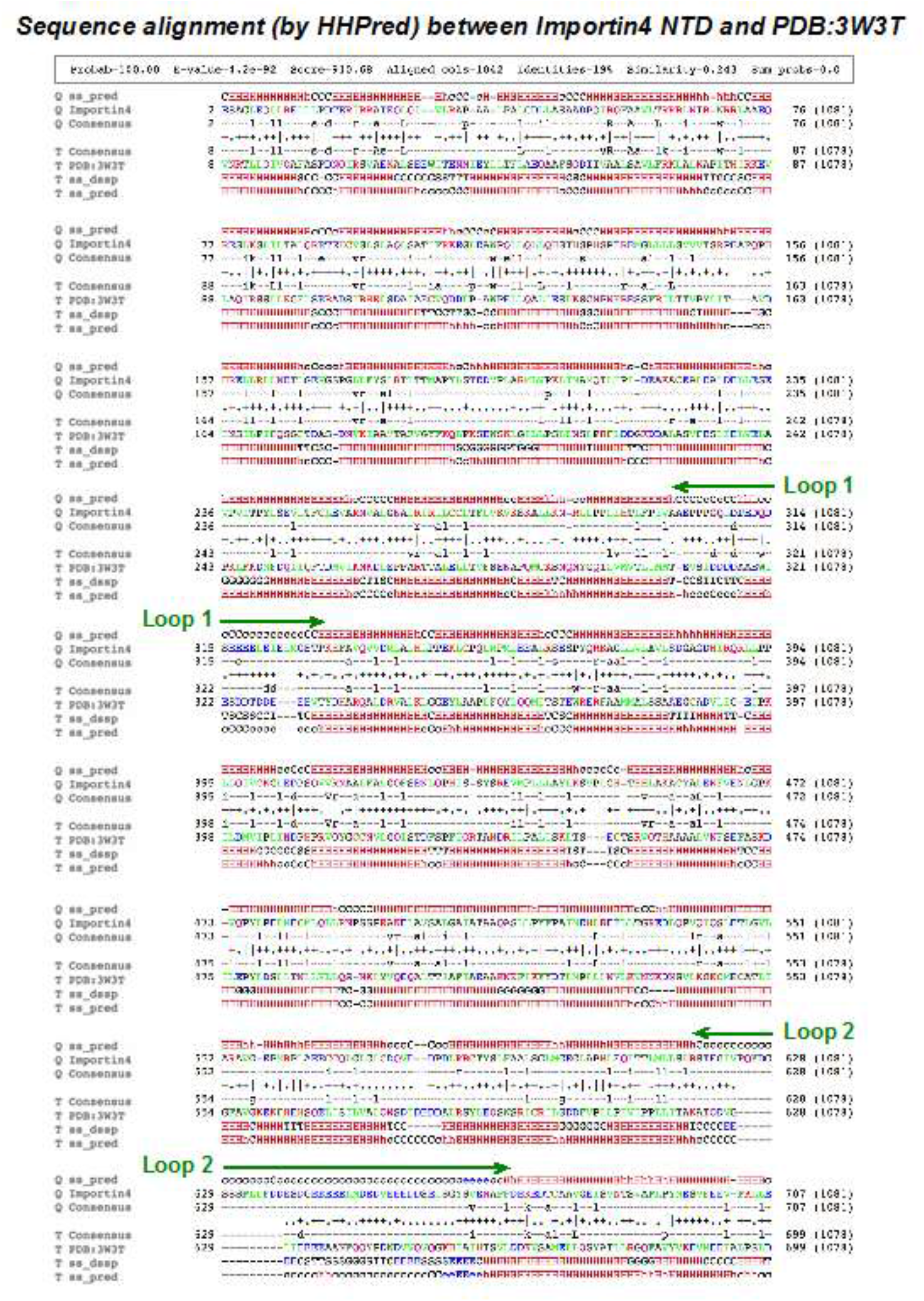
Sequence alignment between the human Importin4 N-terminal domain (NTD) and yeast Kap121p. The sequence alignment between the modeled N-terminal region of the human Importin4 and the template structure of *S.c.* Kap121p is shown, along with the secondary structure assignment by HHPred. The positions of two long acidic loops in the NTD of the human Importin4 are indicated by green arrows.

**Supplementary Fig. 7.**
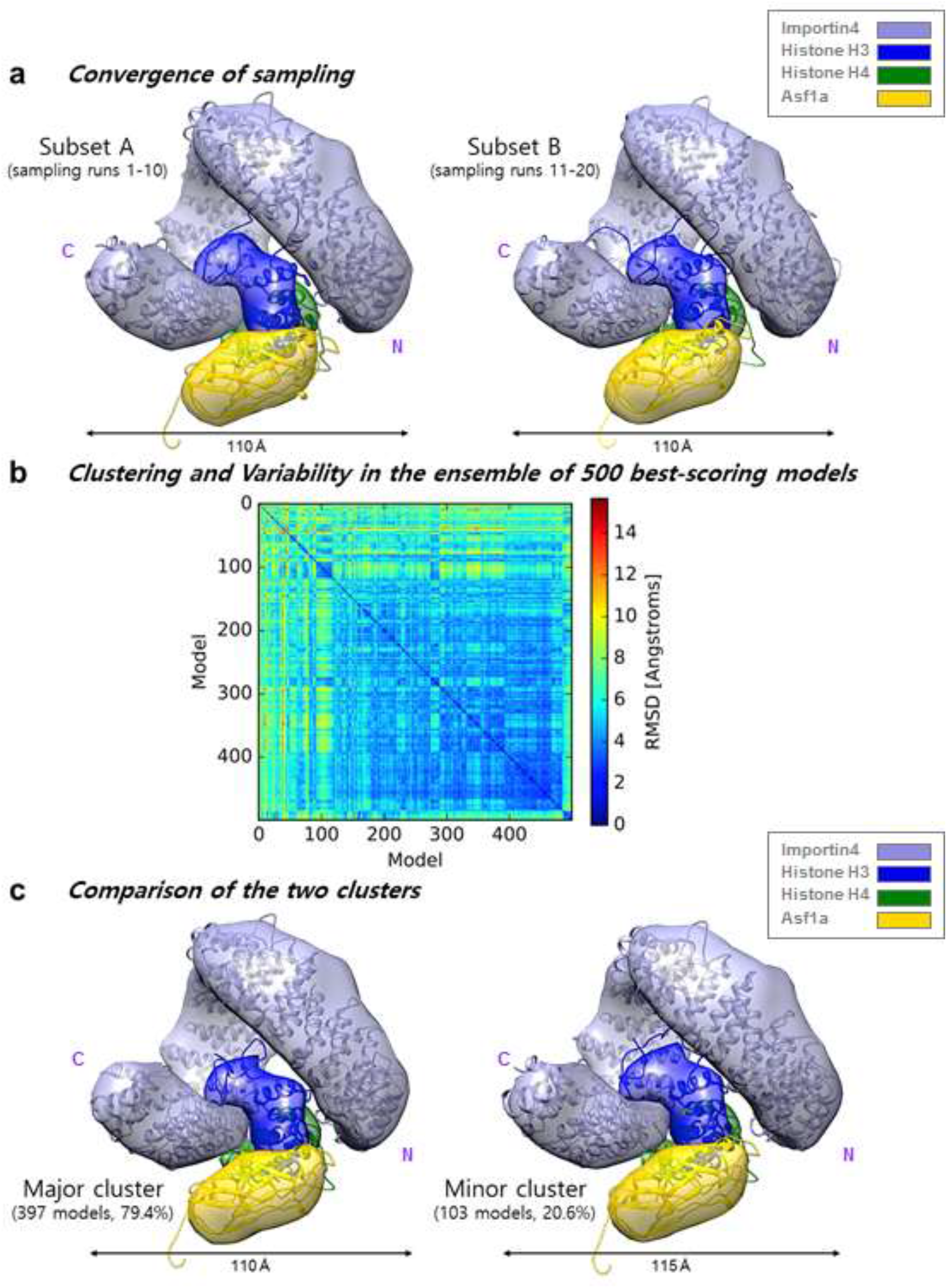
Analysis of the ensemble of 500 best-scoring complex models. (a) Comparison of a subset of 250 best-scoring models from runs 1–10 (left) to another subset of 250 best scoring models from runs 11–20 (right). Each subset of the best-scoring models was converted into a density map that specified how often grid points of the map were occupied by a given protein (the “localization probability density map”). The two localization probability density maps were nearly identical to each other (cross-correlation coefficient = 0.99). Consistently, the TM-score, a measure of structural similarity, between representative (centroid) models of the two subsets is 0.951, indicating about the same fold. (b) The root-mean-square deviation (RMSD) values between a pair of the 500 models is shown. The largest RMSD was ~15 Å. However, most of the RMSD values were less than ~11 Å. (c) Comparison of the two clusters identified by K-means clustering (Shi et al., 2014). The localization of all protein components is nearly identical between the dominant and minor clusters (cross-correlation coefficient = 0.99). The TM-score between representative (centroid) models of the two clusters is 0.889, indicating about the same fold. Therefore, there is effectively a single cluster of the 500 best-scoring models at the model precision of ~11 Å.

**Supplementary Fig. 8.**
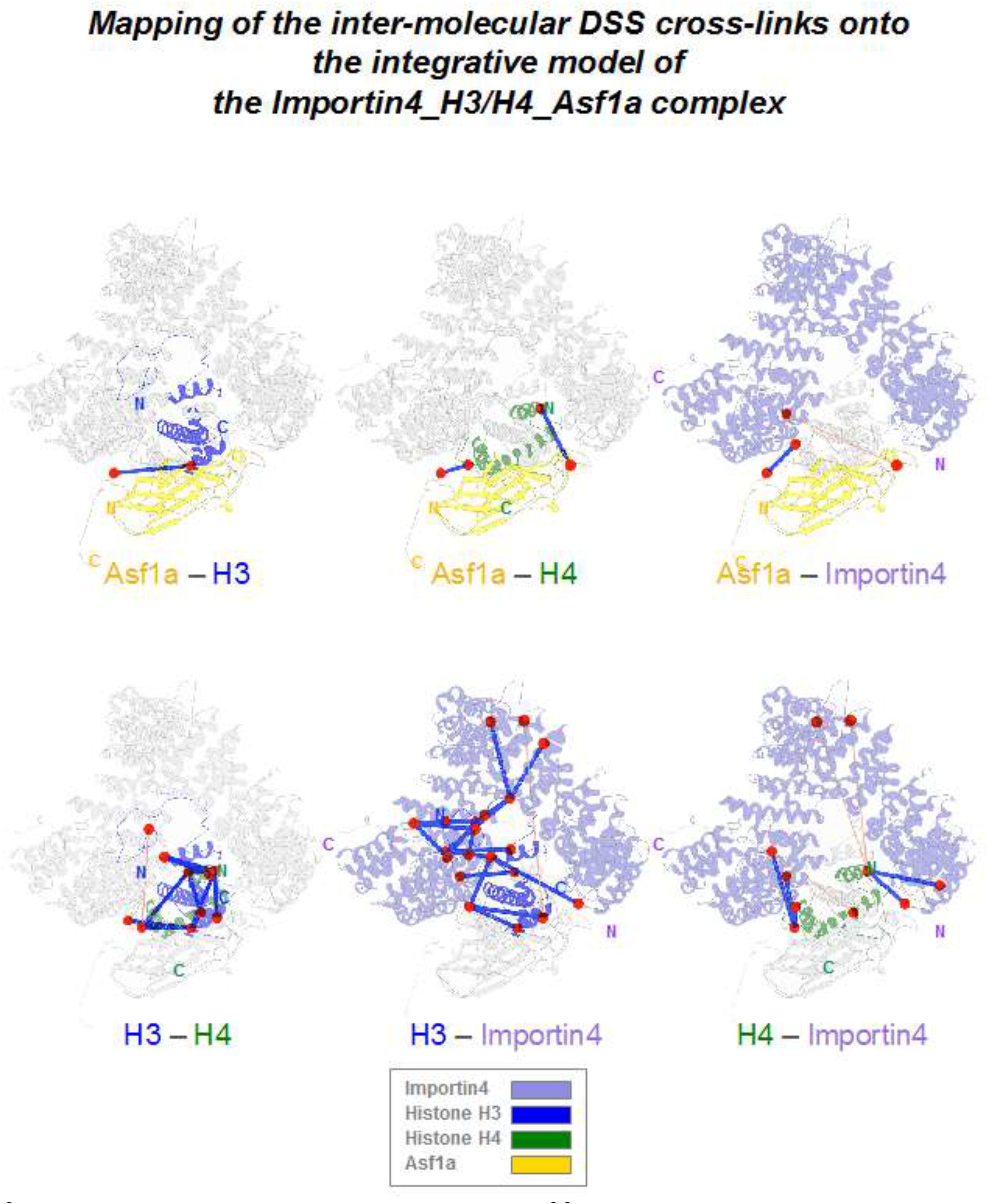
Mapping of the inter-molecular DSS cross-links onto the integrative model of the Importin4_H3/H4_Asf1a complex. Mapping of the inter-molecular cross-links onto the representative model is shown for different proteins. The chemical cross-links falling within the C_α_-C_α_ distance threshold of 35 Å (blue lines) or outside of that threshold (orange lines) are shown on the representative model of the complex (ribbon representation). Each of the crosslinked lysine residues is depicted as a red circle.

**Supplementary Fig. 9.**
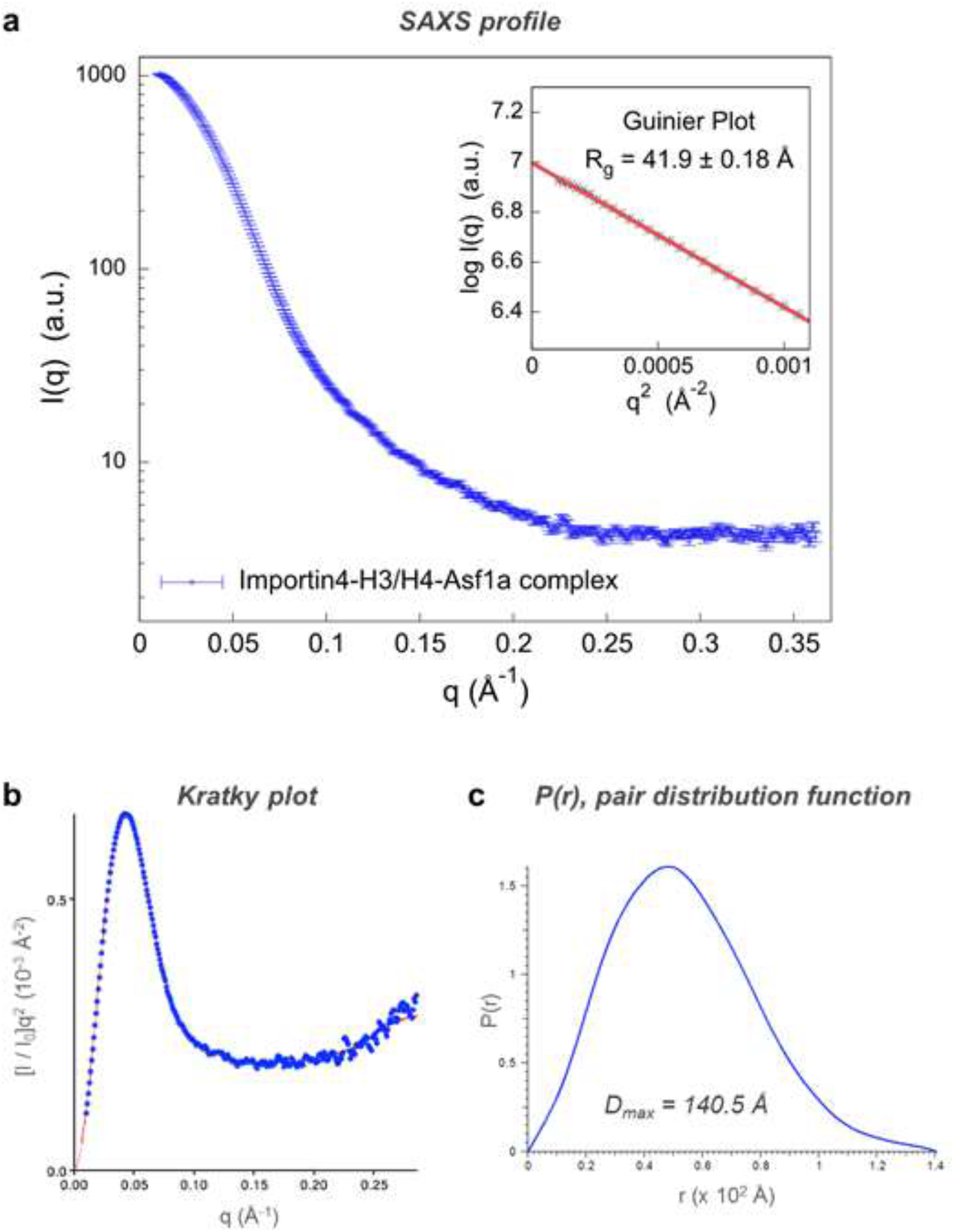
SAXS analysis of Importin4_Histone H3/H4_Asf1a complex. (a) The SAXS profile of the Importin4_histone H3/H4_Asf1a complex is shown in blue dots, along with the extrapolation curve (red). The molecular weight of the SAXS sample was estimated to be 162.1 kDa by using SAXS MOW, a value nearly identical to the expected molecular weight of 161 kDa from the sequence (thus confirming the monomer state of the SAXS sample). An inset presents the Guinier plot shown as a red line in the Guinier region (points 3 to 21). The linearity in the Guinier plot confirms a high degree of compositional homogeneity of the complex in solution. The radius of gyration (*R_g_*) was determined as 41.9 ± 0.2 Å. (b) The Kratky plot is shown in blue dots, along with the extrapolation curve (red). A plateau in this plot (q=0.15 - 0.25 Å^-1^) suggests some flexibility in the complex. (c) The pair distribution function (P(r)) is shown. The maximum particle size (*D_max_*) was determined as 140.5 Å

**Supplementary Fig. 10.**
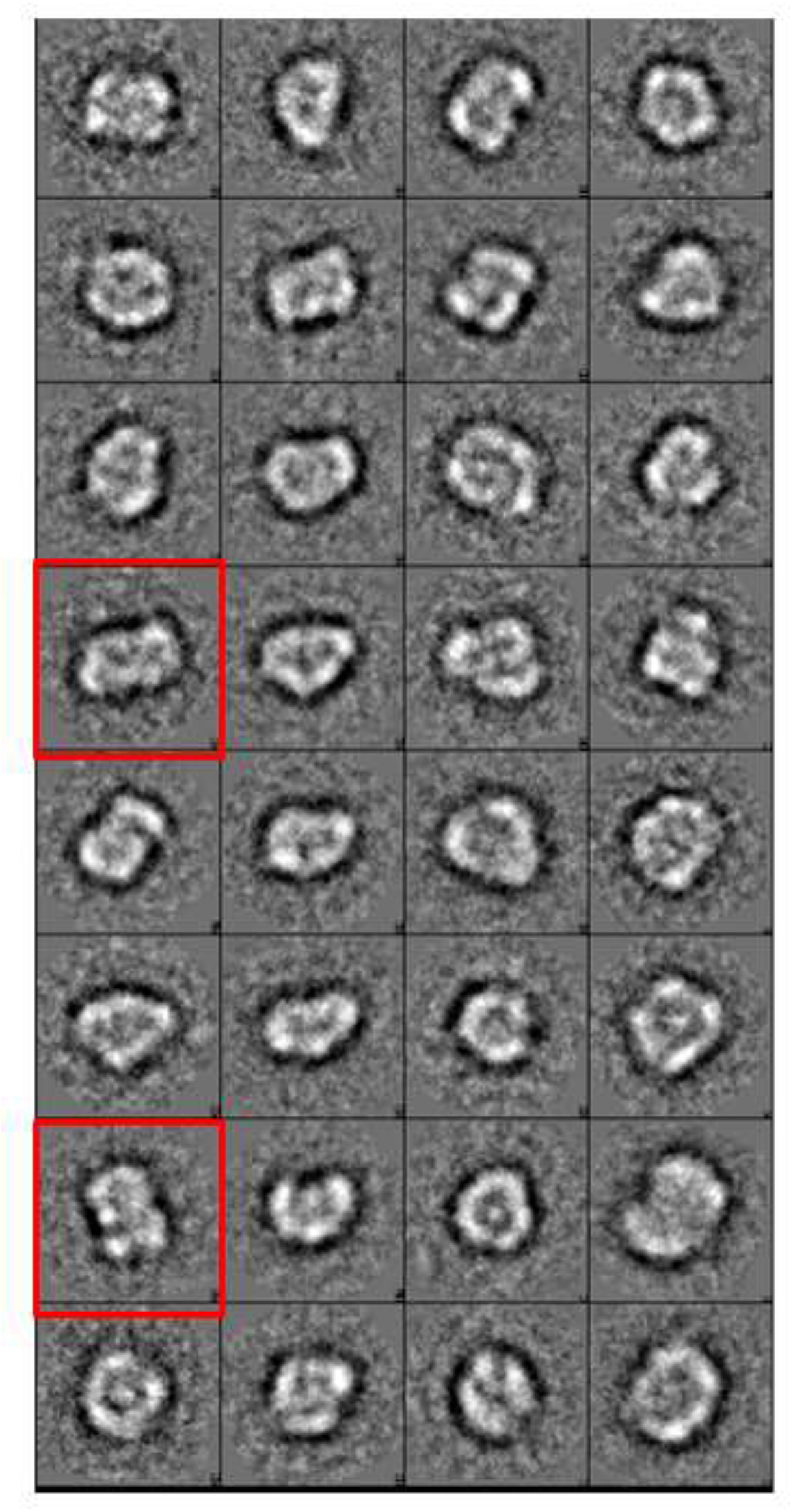
Negative-stain EM 2D class-averages of Importin4_Histone H3/H4_Asf1a complex. 2D class-averages of Importin4_Histone H3/H4_Asf1a complex by negative-stain EM analysis. The extended conformations of the complex observed in the 2D class-average are boxed in red.

**Supplementary Table 1.** List of the positions of the inter- and intra-molecular crosslinks

| Intermolecular cross-links |  |  |  |  |
| --- | --- | --- | --- | --- |
| Protein 1 | Protein 2 | Residue 1 | Residue 2 | Cα-Cα<br>Distance in<br>the model (Å) |
| Importin4 | Asf1a | 867 | 159 | 15.9 |
| Importin4 | Asf1a | 801 | 129 | 58.5 |
| Importin4 | H3 | 110 | 56 | 34.8 |
| Importin4 | H3 | 370 | 27 | 33.0 |
| Importin4 | H3 | 401 | 122 | 67.7 |
| Importin4 | H3 | 497 | 27 | 27.7 |
| Importin4 | H3 | 715 | 4 | 20.9 |
| Importin4 | H3 | 715 | 9 | 20.1 |
| Importin4 | H3 | 715 | 14 | 27.8 |
| Importin4 | H3 | 715 | 18 | 21.9 |
| Importin4 | H3 | 715 | 23 | 11.2 |
| Importin4 | H3 | 715 | 27 | 21.4 |
| Importin4 | H3 | 788 | 27 | 33.3 |
| Importin4 | H3 | 788 | 56 | 28.3 |
| Importin4 | H3 | 801 | 64 | 21.2 |
| Importin4 | H3 | 788 | 36 | 22.8 |
| Importin4 | H3 | 867 | 56 | 21.3 |
| Importin4 | H3 | 867 | 79 | 27.2 |
| Importin4 | H3 | 867 | 122 | 29.2 |
| Importin4 | H3 | 1035 | 4 | 23.0 |
| Importin4 | H3 | 1035 | 14 | 18.4 |
| Importin4 | H3 | 1035 | 18 | 15.9 |
| Importin4 | H4 | 81 | 31 | 25.9 |
| Importin4 | H4 | 110 | 31 | 16.5 |
| Importin4 | H4 | 401 | 31 | 60.5 |
| Importin4 | H4 | 497 | 31 | 55.6 |
| Importin4 | H4 | 788 | 77 | 26.3 |
| Importin4 | H4 | 801 | 77 | 18.0 |
| Importin4 | H4 | 788 | 59 | 36.6 |
| Importin4 | H4 | 867 | 77 | 16.4 |
| Asf1a | H3 | 159 | 79 | 29.0 |
| Asf1a | H4 | 159 | 77 | 12.6 |
| Asf1a | H4 | 129 | 31 | 30.0 |
| H4 | H3 | 44 | 56 | 16.2 |
| H4 | H3 | 31 | 56 | 20.3 |
| H4 | H3 | 31 | 64 | 14.4 |
| H4 | H3 | 31 | 79 | 29.1 |
| H4 | H3 | 31 | 122 | 18.7 |
| H4 | H3 | 59 | 64 | 15.7 |
| H4 | H3 | 59 | 79 | 16.7 |
| H4 | H3 | 77 | 4 | 39.3 |
| H4 | H3 | 77 | 64 | 24.2 |
| H4 | H3 | 77 | 79 | 17.9 |
| H4 | H3 | 79 | 79 | 23.4 |

| Intramolecular cross-links |  |  |  |  |
| --- | --- | --- | --- | --- |
| Protein 1 | Protein 2 | Residue 1 | Residue 2 | Distance (Å) |
| Importin4 | Importin4 | 715 | 788 | 15.3 |
| Importin4 | Importin4 | 788 | 867 | 24.5 |
| Importin4 | Importin4 | 801 | 788 | 8.5 |
| Importin4 | Importin4 | 801 | 867 | 19.7 |
| Importin4 | Importin4 | 940 | 867 | 21.8 |
| H3 | H3 | 27 | 23 | 12.7 |
| H3 | H3 | 27 | 56 | 21.6 |
| H3 | H3 | 36 | 56 | 13.1 |
| H3 | H3 | 36 | 64 | 11.7 |
| H3 | H3 | 37 | 56 | 13.9 |
| H3 | H3 | 37 | 64 | 9.6 |
| H3 | H3 | 37 | 79 | 31.1 |
| H3 | H3 | 56 | 4 | 10.6 |
| H3 | H3 | 56 | 14 | 15.8 |
| H3 | H3 | 56 | 18 | 22.8 |
| H3 | H3 | 56 | 23 | 17.4 |
| H3 | H3 | 56 | 64 | 20.1 |
| H3 | H3 | 79 | 4 | 45.6 |
| H3 | H3 | 79 | 56 | 39.5 |
| H3 | H3 | 79 | 64 | 22.5 |
| H3 | H3 | 122 | 64 | 28.2 |
| H3 | H3 | 122 | 79 | 33.0 |
| H4 | H4 | 31 | 44 | 15.5 |
| H4 | H4 | 31 | 59 | 15.0 |
| H4 | H4 | 31 | 77 | 34.9 |
| H4 | H4 | 59 | 8 | 28.9 |
| H4 | H4 | 77 | 8 | 12.3 |
| H4 | H4 | 79 | 77 | 6.2 |

## References

[1] Luger K, Mader AW, Richmond RK, Sargent DF, Richmond TJ. Crystal structure of the nucleosome core particle at 2.8[thinsp]A resolution. Nature. 1997;389:251–60.

[2] Venkatesh S, Workman JL. Histone exchange, chromatin structure and the regulation of transcription. Nat Rev Mol Cell Biol. 2015;16:178–89.

[3] Tagami H, Ray-Gallet D, Almouzni G, Nakatani Y. Histone H3.1 and H3.3 Complexes Mediate Nucleosome Assembly Pathways Dependent or Independent of DNA. Synthesis. 2004;116:51–61.

[4] Campos EI, Fillingham J, Li G, Zheng H, Voigt P, Kuo WH, et al. The program for processing newly synthesized histones H3.1 and H4. Nat Struct Mol Biol. 2010;17:1343–51.

[5] Hammond CM, Stromme CB, Huang H, Patel DJ, Groth A. Histone chaperone networks shaping chromatin function. Nat Rev Mol Cell Biol. 2017;18:141–58.

[6] Smith S, Stillman B. Purification and characterization of CAF-I, a human cell factor required for chromatin assembly during DNA replication in vitro. Cell. 1989;58:15–25.

[7] Natsume R, Eitoku M, Akai Y, Sano N, Horikoshi M, Senda T. Structure and function of the histone chaperone CIA/ASF1 complexed with histones H3 and H4. Nature. 2007;446:338–41.

[8] English CM, Adkins MW, Carson JJ, Churchill ME, Tyler JK. Structural basis for the histone chaperone activity of Asf1. Cell. 2006;127:495–508.

[9] Park Y-J, Luger K. The structure of nucleosome assembly protein 1. Proceedings of the National Academy of Sciences. 2006;103:1248–53.

[10] Mosammaparast N, Guo Y, Shabanowitz J, Hunt DF, Pemberton LF. Pathways mediating the nuclear import of histones H3 and H4 in yeast. J Biol Chem. 2002;277:862–8.

[11] Fukuhara N, Fernandez E, Ebert J, Conti E, Svergun D. Conformational variability of nucleo-cytoplasmic transport factors. J Biol Chem. 2004;279:2176–81.

[12] Görlich D KU. Transport Between the Cell Nucleus and the Cytoplasm. Annual Review of Cell and Developmental Biology. 1999;15:607–60.

[13] Lounsbury KM, Macara IG. Ran-binding Protein 1 (RanBP1) Forms a Ternary Complex with Ran and Karyopherin β and Reduces Ran GTPase-activating Protein (RanGAP) Inh ibition by Karyopherin β. Journal of Biological Chemistry. 1997;272:551–5.

[14] Jäkel S, Mingot JM, Schwarzmaier P, Hartmann E, Görlich D. Importins fulfil a dual function as nuclear import receptors and cytoplasmic chaperones for exposed basic domains. The EMBO Journal. 2002;21:377–86.

[15] Miyauchi Y, Michigami T, Sakaguchi N, Sekimoto T, Yoneda Y, Pike JW, et al. Importin 4 is responsible for ligand-independent nuclear translocation of vitamin D receptor. J Biol Chem. 2005;280:40901–8.

[16] Lokich E, Singh RK, Han A, Romano N, Yano N, Kim K, et al. HE4 expression is associated with hormonal elements and mediated by importin-dependent nuclear translocation. Sci Rep. 2014;4:5500.

[17] Alvarez F, Munoz F, Schilcher P, Imhof A, Almouzni G, Loyola A. Sequential establishment of marks on soluble histones H3 and H4. J Biol Chem. 2011;286:17714–21.

[18] Baake M, Doenecke D, Albig W. Characterisation of nuclear localisation signals of the four human core histones. J Cell Biochem. 2001;81:333–46.

[19] Blackwell JS, Jr., Wilkinson ST, Mosammaparast N, Pemberton LF. Mutational analysis of H3 and H4 N termini reveals distinct roles in nuclear import. J Biol Chem. 2007;282:20142–50.

[20] Greiner M, Caesar S, Schlenstedt G. The histones H2A/H2B and H3/H4 are imported into the yeast nucleus by different mechanisms. Eur J Cell Biol. 2004;83:511–20.

[21] Johnson-Saliba M, Siddon NA, Clarkson MJ, Tremethick DJ, Jans DA. Distinct importin recognition properties of histones and chromatin assembly factors. FEBS Lett. 2000;467:169–74.

[22] Muhlhausser P, Muller EC, Otto A, Kutay U. Multiple pathways contribute to nuclear import of core histones. EMBO Rep. 2001;2:690–6.

[23] Soniat M, Cagatay T, Chook YM. Recognition Elements in the Histone H3 and H4 Tails for Seven Different Importins. J Biol Chem. 2016;291:21171–83.

[24] Soniat M, Chook YM. Karyopherin-beta2 Recognition of a PY-NLS Variant that Lacks the Proline-Tyrosine Motif. Structure. 2016;24:1802–9.

[25] Opel M, Lando D, Bonilla C, Trewick SC, Boukaba A, Walfridsson J, et al. Genome-wide studies of histone demethylation catalysed by the fission yeast homologues of mammalian LSD1. PLoS One. 2007;2:e386.

[26] Groves MR, Hanlon N, Turowski P, Hemmings BA, Barford D. The Structure of the Protein Phosphatase 2A PR65/A Subunit Reveals the Conformation of Its 15 Tandemly Repeated HEAT Motifs. Cell. 1999;96:99–110.

[27] Cho US, Xu W. Crystal structure of a protein phosphatase 2A heterotrimeric holoenzyme. Nature. 2007;445:53–7.

[28] Kobayashi J, Matsuura Y. Structural Basis for Cell-Cycle-Dependent Nuclear Import Mediated by the Karyopherin Kap121p. Journal of Molecular Biology. 2013;425:1852–68.

[29] Russel D, Lasker K, Webb B, Velazquez-Muriel J, Tjioe E, Schneidman-Duhovny D, et al. Putting the pieces together: integrative modeling platform software for structure determination of macromolecular assemblies. PLoS Biol. 2012;10:e1001244.

[30] Sali A, Blundell TL. Comparative protein modelling by satisfaction of spatial restraints. J Mol Biol. 1993;234:779–815.

[31] Soding J. Protein homology detection by HMM-HMM comparison. Bioinformatics. 2005;21:951–60.

[32] Soding J, Biegert A, Lupas AN. The HHpred interactive server for protein homology detection and structure prediction. Nucleic Acids Res. 2005;33:W244–8.

[33] Fischer H, Neto MD, Napolitano HB, Polikarpov I, Craievich AF. Determination of the molecular weight of proteins in solution from a single small-angle X-ray scattering measurement on a relative scale. Journal of Applied Crystallography. 2010;43:101–9.

[34] Alabert C, Groth A. Chromatin replication and epigenome maintenance. Nat Rev Mol Cell Biol. 2012;13:153–67.

[35] Annunziato AT. Assembling chromatin: The long and winding road. Biochimica et Biophysica Acta (BBA) - Gene Regulatory Mechanisms. 2012;1819:196–210.

[36] Clemente-Ruiz M, Gonzalez-Prieto R, Prado F. Histone H3K56 acetylation, CAF1, and Rtt106 coordinate nucleosome assembly and stability of advancing replication forks. PLoS Genet. 2011;7:e1002376.

[37] Krawitz DC, Kama T, Kaufman PD. Chromatin assembly factor I mutants defective for PCNA binding require Asf1/Hir proteins for silencing. Mol Cell Biol. 2002;22:614–25.

[38] Kim D, Setiaputra D, Jung T, Chung J, Leitner A, Yoon J, et al. Molecular Architecture of Yeast Chromatin Assembly Factor 1. Scientific Reports. 2016;6:26702.

[39] Campos EI, Smits AH, Kang YH, Landry S, Escobar TM, Nayak S, et al. Analysis of the Histone H3.1 Interactome: A Suitable Chaperone for the Right. Event. 2015;60:697–709.

[40] Green EM, Antczak AJ, Bailey AO, Franco AA, Wu KJ, Yates JR, 3rd, et al. Replication-independent histone deposition by the HIR complex and Asf1. Curr Biol. 2005;15:2044–9.

[41] Forwood JK, Lange A, Zachariae U, Marfori M, Preast C, Grubmuller H, et al. Quantitative structural analysis of importin-beta flexibility: paradigm for solenoid protein structures. Structure. 2010;18:1171–83.

[42] Leitner A, Walzthoeni T, Aebersold R. Lysine-specific chemical cross-linking of protein complexes and identification of cross-linking sites using LC-MS/MS and the xQuest/xProphet software pipeline. Nat Protoc. 2014;9:120–37.

[43] Walzthoeni T, Claassen M, Leitner A, Herzog F, Bohn S, Forster F, et al. False discovery rate estimation for cross-linked peptides identified by mass spectrometry. Nat Methods. 2012;9:901–3.

[44] Otwinowski Z, Minor W. Processing of X-ray diffraction data collected in oscillation mode. Methods Enzymol. 1997;276:307–26.

[45] McCoy AJ, Grosse-Kunstleve RW, Adams PD, Winn MD, Storoni LC, Read RJ. Phaser crystallographic software. J Appl Crystallogr. 2007;40:658–74.

[46] Brunger AT. Version 1.2 of the Crystallography and NMR. system. 2007;2:2728–33.

[47] Emsley P, Lohkamp B, Scott WG, Cowtan K. Features and development of Coot. Acta Crystallogr D Biol Crystallogr. 2010;66:486–501.

[48] Tang G, Peng L, Baldwin PR, Mann DS, Jiang W, Rees I, et al. EMAN2: an extensible image processing suite for electron. microscopy. 2007;157:38–46.

[49] Alber F, Dokudovskaya S, Veenhoff LM, Zhang W, Kipper J, Devos D, et al. Determining the architectures of macromolecular assemblies. Nature. 2007;450:683–94.

[50] Shi Y, Fernandez-Martinez* J, Tjioe* E, Pellarin* R, Kim* SJ, Williams R, et al. Structural characterization by cross-linking reveals the detailed architecture of a coatomer-related heptameric module from the nuclear pore complex. Mol Cell Proteomics. 2014;13:2927–43.

[51] Fernandez-Martinez J, Kim SJ, Shi Y, Upla P, Pellarin R, Gagnon M, et al. Structure and Function of the Nuclear Pore Complex Cytoplasmic mRNA Export Platform. Cell. 2016;167:1215–28e25.

[52] Shen MY, Sali A. Statistical potential for assessment and prediction of protein structures. Protein Sci. 2006;15:2507–24.

[53] Cingolani G, Lashuel HA, Gerace L, Muller CW. Nuclear import factors importin alpha and importin beta undergo mutually induced conformational changes upon association. FEBS Lett. 2000;484:291–8.

[54] Imasaki T, Shimizu T, Hashimoto H, Hidaka Y, Kose S, Imamoto N, et al. Structural basis for substrate recognition and dissociation by human transportin 1. Mol Cell. 2007;28:57–67.

[55] Emekli U, Schneidman-Duhovny D, Wolfson HJ, Nussinov R, Haliloglu T. HingeProt: automated prediction of hinges in protein structures. Proteins. 2008;70:1219–27.

[56] LoPiccolo J, Kim SJ, Shi Y, Wu B, Wu H, Chait BT, et al. Assembly and Molecular Architecture of the Phosphoinositide 3-Kinase p85alpha Homodimer. J Biol Chem. 2015;290:30390–405.

[57] Zhang Y, Skolnick J. Scoring function for automated assessment of protein structure template quality. Proteins. 2004;57:702–10.

